# Stress- and metabolic responses of *Candida albicans* require Tor1 kinase N-terminal HEAT repeats

**DOI:** 10.1101/2021.11.05.467412

**Authors:** Wanjun Qi, Maikel Acosta-Zaldivar, Peter R. Flanagan, Ning-Ning Liu, Niketa Jani, José F. Fierro, María T. Andrés, Gary P. Moran, Julia R. Köhler

## Abstract

Whether to commit limited cellular resources toward growth and proliferation, or toward survival and stress responses, is an essential determination made by Target of Rapamycin Complex 1 (TORC1) for a eukaryotic cell in response to favorable or adverse conditions. Loss of TORC1 function is lethal. The TORC1 inhibitor rapamycin that targets the highly conserved Tor kinase domain kills fungal pathogens like *Candida albicans*, but is also severely toxic to human cells. The least conserved region of fungal and human Tor kinases are the N-terminal HEAT domains. We examined the role of the 8 most N-terminal HEAT repeats of *C. albicans* Tor1. We compared nutritional- and stress responses of cells expressing a message for N-terminally truncated Tor1 from repressible *tetO*, with cells expressing a wild type *TOR1* allele from *tetO* or from the native promoter. Some but not all stress responses were significantly impaired by loss of Tor1 N-terminal HEAT repeats, including those to oxidative-, cell wall-, and heat stress; in contrast, plasma membrane stress and antifungal agents that disrupt plasma membrane function were tolerated by cells lacking this Tor1 region. Translation was inappropriately upregulated during oxidative stress in cells lacking N-terminal Tor1 HEAT repeats despite simultaneously elevated Gcn2 activity, while activation of the oxidative stress response MAP kinase Hog1 was weak. Conversely, these cells were unable to take advantage of favorable nutritional conditions by accelerating their growth. While consuming oxygen more slowly than cells containing wild type *TOR1* alleles during growth in glucose, cells lacking N-terminal Tor1 HEAT repeats were capable of utilizing non-fermentable as well as fermentable carbon sources and of growth during severe acid stress, but were uniquely unable to grow on lactate. Genome-wide expression analysis showed paradoxical simultaneous activation of anabolic- and starvation responses in cells lacking Tor1 N-terminal HEAT repeats, with misregulation of carbon metabolism and of translational machinery biosynthesis. Targeting fungal-specific Tor1 N-terminal HEAT repeats with small molecules might abrogate fungal viability, especially when during infection multiple stresses are imposed by the host immune system.

## Introduction

The Target of Rapamycin Complex 1 (TORC1) makes fundamental decisions in the life of a eukaryotic cell. It collects information from numerous sources on conditions that affect the cell’s chances of successful growth and proliferation. It then directs downstream regulators to promote anabolic- or stress- and survival responses. Called “the brain of the cell” by Michael Hall [1], TORC1, by extension of this metaphor, integrates data from a plethora of afferent projections in order to send distinct efferent signals to its effectors. In contrast to the brain, in which afferent and efferent fibers can be anatomically separated, TORC1 has only its enormous protein-protein interaction domains to invite upstream signaling molecules to dock and to select its downstream targets for phosphorylation, or to exclude them as appropriate from the kinase catalytic domain. Considering the large number of signaling inputs into TORC1 and its substantial number of effectors, very little is known about where the relevant afferent and efferent components of the pathway interact with components of the complex.

Appropriate responses to changing favorable or unfavorable environmental conditions are crucial for all free-living microorganisms. Saprophytes like *Saccharomyces cerevisiae* experience fluctuations of nutrient availability, temperature, osmolarity, oxygen tension and redox potential. Controlled cessation of anabolic activity and regulated exit from the cell cycle as well as induction of stress responses promote survival during adverse conditions, while prompt resumption of translation, organelle biosynthesis and DNA replication enhance competitive fitness in a favorable environment. A commensal and pathogen like *Candida albicans* additionally contends with the density of competing flora in the host gastrointestinal tract, and with stresses actively imposed by the host immune system. These stresses include membrane stress induced by antimicrobial peptides, oxidative- and cell wall stress generated by host phagocytes and heat stress through fever as a host defense.

TORC1 was discovered through genetic analysis of the mechanism of action of rapamycin [2], a streptomycetal secondary metabolite originally isolated for its antifungal activity against *C. albicans* [3, 4]. Found to be toxic for human cells, especially for lymphocytes whose protein production and proliferation are integral to their function, rapamycin’s development as an antifungal agent was abandoned [5]. The discovery of its target over a decade later [2] set in motion a plethora of novel areas of investigation now known to be central to eukaryotic cell biology. Given rapamycin’s cidal potency against *C. albicans* and other fungi [6, 7], as well as against other pathogens relying on eukaryotic mechanisms of growth and proliferation control [8], inhibitors with specificity for the fungal- and neutrality toward the human TORC1 could be of great therapeutic interest [6]. In order to develop such compounds, more detailed understanding of the functional role of each of the Tor kinase protein domains and their subsegments, as well as of the structural differences between fungal and mammalian TORC1, will be a precondition.

In *S. cerevisiae*, TORC1 is a dimer of each of the components, the kinases Tor1 or Tor2, the TORC1-defining component Kontroller of Growth (Kog1), the WD40 repeat-containing regulator and stabilizer of the TORC1 complex, Lethal with Sec13 protein 8 (Lst8) [9], and Tor Complex One 89 kD protein (Tco89), noted for its role in cell wall integrity [10] [11]. *C. albicans*, like mammals, has only one Tor kinase, Tor1. Homologs of the other *S. cerevisiae* TORC1 components are also present in *C. albicans*.

Tor kinase is a serine/threonine protein kinase despite its homology to phosphatidylinositol 3-kinases. Its domain structure is conserved to a large extent across eukaryotes, so that fungal, mammalian and plant structures have been used to analyze the functional units [12]. Its C-terminal kinase domain consists of an N-terminal and a C-terminal lobe that together form a deep catalytic cleft which enables inhibition by blocking substrate access [13]. At the N-terminus of Tor kinase are two domains of helical repeats, the N-terminal and the middle HEAT (huntingtin, elongation factor 3, a subunit of PP2A, and TOR1) domains, followed by the FAT (named for FKBP12 Rapamycin Associated Protein (FRAP), ATM, TRRAP) domain. The FRB (FKBP12-Rapamycin binding) domain is part of the N-terminal lobe of the kinase domain, and in mTOR acts as a gatekeeper to control substrate access to the catalytic site [13]. At its C-terminus the kinase domain is flanked by C-terminal FAT domains [14].

N terminal HEAT repeats [15] consist of arrays of two antiparallel alpha helices of 10-20 amino acid residues’ length each separated by intraunit loops of 5-8 residues [15]. *S. cerevisiae* TOR kinase N-terminal HEAT repeats were shown by electron microscopy to interact closely with the C-terminus of Kog1 [16], a scaffold for interacting proteins and a gatekeeper for substrates to the kinase active site [17]. A cryo-EM structure of Tor kinase complexed with Lst8 of the yeast *Kluyveromyces marxianus* showed its N terminus to consist of a curved “spiral” of 36 helices comprising about 800 residues, representing the N-terminal HEAT domain, followed by a linker that connects to another “bridge” sequence of 14 helices comprising ∼400 residues, representing the middle HEAT domain [18]. The major interaction sites between two KmTor kinase molecules in the dyadic complex described by these authors are formed reciprocally between the N-terminal (spiral) and middle (bridge) HEAT domains of the two Tor kinase molecules which connect in a yin-yang orientation. In their projection, Kog1 binds to the N-terminal and middle HEAT domain of the opposite Tor kinase subunit in the complex, contributing to the stability of the dimer [18].

The N-terminal HEAT domain of mammalian Tor kinase is predicted to be exposed at the surface of mTORC1 and bind its regulators [17]. As deduced from aligning the CaTor1 sequence with mTOR, whose HEAT repeats include the N-terminal 1382 amino acid residues [19], CaTor1 HEAT repeats comprise 1334 N-terminal amino acids. N-terminal truncation of mTOR by 297 residues weakens its interaction with the conserved Kog1 homolog Raptor [20].

An N-terminally truncated human mTOR consisting only of residues 1376 to 2549, in complex with full-length human mLST8, was able to phosphorylate a classic TORC1 substrate, S6 kinase1 (S6K1), in vitro [13], highlighting that Tor kinase HEAT repeats and Kog1 are not necessary for catalytic activity; they serve to recruit substrates for, and to regulate the kinase. Recent structural analysis of mTOR has shown that binding of its activator Rheb induces conformational changes involving the N-HEAT domain and the FAT domain that realign active-site residues in the catalytic cleft to facilitate the enzymatic reaction [12]. It is unclear whether this allosteric regulatory mechanism is conserved in model fungi. In *S. cerevisiae* the Rheb homolog Rhb1 acts on arginine uptake and is not essential [21] whereas it is essential and acts in both basic amino acid uptake and TORC1-controlled processes like growth, differentiation for mating and cell cycle exit, in *Schizosaccharomyces pombe* [22]. In *C. albicans*, Rhb1 and its upstream regulator Tsc2 are involved in rapamycin susceptibility and regulate filamentous growth and expression of the ammonium transporter Mep2 and the secreted proteinase Sap2 [23, 24]. CaRhb1 is required for normal cell wall stress resistance and cell wall integrity signaling [23]. It is required for appropriate Rps6 phosphorylation, a downstream effect of active TORC1 [25]; deletion mutants in *RHB1* differentially transcribe genes controlled by TORC1 like those involved in ribosome biogenesis and amino acid biosynthesis [24, 26]. Rhb1 hence is a component of the *C. albicans* TORC1 signaling pathway but details of its physical interaction with Tor1, if any, are unknown.

Inhibition of mTor kinase by PRAS40 involves its binding to the FRB domain and obstructing the catalytic cleft [12]. This regulatory mechanism is not present in yeast or *C. albicans* which do not have a PRAS40 homolog. Other structural analyses of mTor kinase, including its interactions with the major substrates 4EBP1 and S6K1, similarly cannot be extrapolated to the function of *S. cerevisiae* and *C. albicans* TORC1, since regulating translation initiation does not involve a 4EBP1 homolog in these fungi and the orthologous role of Sch9 as an S6K1 homolog is still debated [27, 28]. Therefore, despite higher-level structural similarities between fungal and mammalian TORC1, Tor kinase activity is regulated differently in fungi than in mammals and the roles of its distinct domains and their physical interaction partners are not understood in detail.

The N-termini of fungal and human Tor kinase are their most widely divergent domains, suggesting they may be sufficiently structurally and functionally distinct that differential chemical targeting could be achievable. The large size and complexity of the N- and M-HEAT domains [18] indicates commensurate complexity of interactions. A functional dissection of their individual roles may aid in further characterizing afferent as well as efferent signal fluxes of Tor1. Responses to nitrogen source quality and quantity and control of protein synthesis are central to TORC1 signaling [29–31], as recently reviewed e.g. in [32]. In *S. cerevisiae* and *C. albicans*, TORC1 responds to carbon source quality and quantity [31] and it controls carbon source-acquisition and -metabolism genes of *S. cerevisiae* [30]. Phosphate repletion is an afferent signal to TORC1 in *S. cerevisiae* and *C. albicans* and TORC1 activity also affects the *C. albicans* phosphate acquisition system, the PHO regulon [33].

As mentioned above, *C. albicans* must adapt to rapidly shifting nutritional states as a commensal in the human gastrointestinal tract, and to stresses imposed not only by competing flora but also by the human immune system during commensalism, mucosal- and invasive infection. To maximize fitness in these quickly changing environments, TORC1 of *C. albicans* must be finely tunable to an array of environmental parameters in widely varying combinations. The challenges of adapting to varying host environments and the importance of Target of Rapamycin complexes in meeting these challenges are exemplified by the sleeping sickness parasite *Trypanosoma brucei*, whose complex life cycle comprising four cell types alternates between its fly vector and its human host [34]: *T. brucei* has four TOR complexes [35, 36]. *C. albicans* TORC1 is involved in hyphal growth and adhesion [37–40], biofilm formation [26], secretion of aspartyl protease [24], and it responds to nitrogen-, carbon-source [25, 41] and phosphate [33] availability. TORC1 is predicted to be required for *C. albicans* fitness in favorable conditions like repletion of distinct nutrients, as well as in manifold stress conditions encountered in the host.

Given the wide array of stimuli to which TORC1 responds and the broad spectrum of responses it controls, we set out to examine the processes regulated by the most N-terminal segment of the N-terminal HEAT domain in *C. albicans*. We constructed two mutant genotypes in which the single remaining *TOR1* alleles were placed under the control of a repressible promoter: wild type full-length *TOR1* or a 5’-truncated *TOR1* allele whose predicted protein product lacks the 8 most N-terminal HEAT repeats. Cells in which these *TOR1* alleles were overexpressed and partially or fully repressed were examined for their responses to nutritional repletion or starvation, and to stresses to which *C. albicans* is exposed in the host.

## Results

### Tor1 N-terminal HEAT repeats were required to accelerate growth in preferred- and to decrease TORC1 signaling in poor nitrogen sources

Aligning *C. albicans* Tor1 and the *K. marxianus* Tor kinase, whose structure has been elucidated in detail [18], with human mTOR, we noted that the most divergent regions of these orthologs are in their N-termini while their catalytic domains are highly conserved (Fig. 1). Tor kinase N termini consist of arrays of HEAT repeats [15]. In order to test the role of *C. albicans* Tor1 and of its N-terminal HEAT repeats in the response to distinct nutrients and stressors, we constructed strains in which transcription of a single *TOR1* allele is controlled by repressible *tetO*. In the absence of the repressing compound doxycycline, transcription from this construct is known to be high [42]. We mutated the second *TOR1* allele in three independent heterozygous deletion mutants in order to detect and avoid artifacts arising from suppressor mutations; strains were constructed to express either full-length Tor1 (Tor1-FL) or an N-terminally truncated Tor1 whose Start methionine is residue 382 of the full length protein (Tor1-Del381). We used the descriptor *TOR1-Del381* for this *TOR1* allele as well, for the sake of simplicity, though the number 381 refers to the truncated amino acid residues and not to the deleted nucleotides. The mutant protein lacks the 8 N-terminal HEAT repeats, truncating the N-terminal HEAT domain, a predicted interaction site with the regulator Kog1. We chose at least 2 strains from each genotype lineage whose phenotypes were indistinguishable for further analysis of growth- and stress phenotypes. We also used two isogenic wild type strains whose deleted *ARG4* locus was independently restored to one wild type allele (resulting in *ARG4/arg4*). To simplify strain descriptors going forward, we note that the designation “Del381” or “FL” refers to strains in which the 5’-truncated *TOR1* allele (encoding an N-terminally truncated protein) or the wild type *TOR1* gene, respectively, were expressed from *tetO*; their respective genotypes were *tor1/tetO-TOR1-Del381* or *tor1/tetO-TOR1*.

**Fig. 1.**
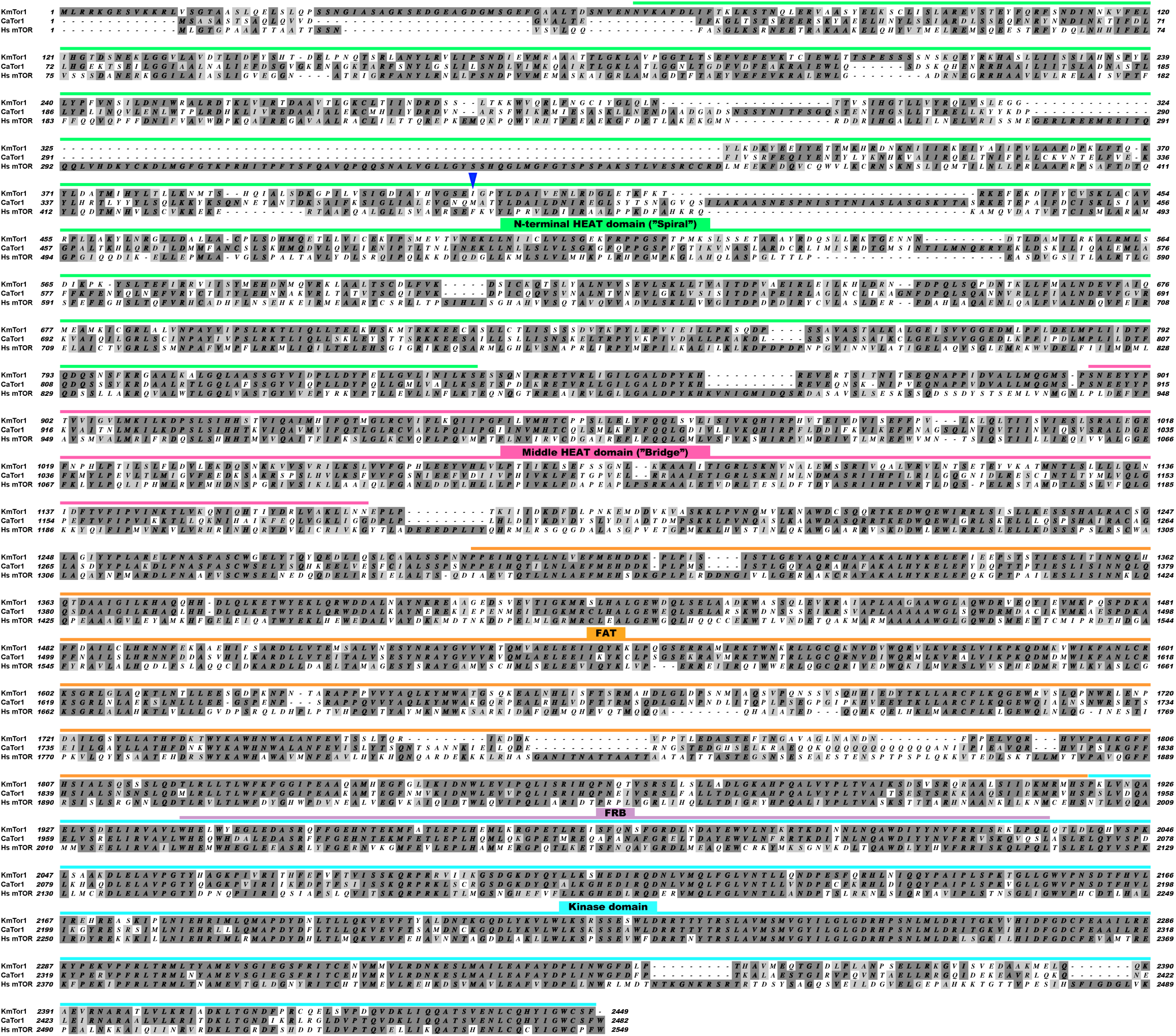
Sequence alignment of fungal and human Tor kinases shows the most divergent regions to lie at the N-terminus. Tor kinase sequences from *Kluyveromyces marxianus* (KmTor1), *Candida albicans* (CaTor1) and human (Hs mTOR) were aligned in MacVector Software using ClustalW (v1.83) multiple sequence alignment. The blue arrow 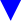 marks the beginning of the Tor1-Del381 sequence. Domain annotation is adapted from Baretic et al. [18] according to the KmTor1 sequence.

Growth of cells expressing full-length or truncated *TOR1* from *tetO* decreased in increasing concentrations of doxycycline, as expected (Fig. 2A). When *TOR1* expression was fully repressed at high concentrations of doxycycline, 1 or 2 µg/ml (Fig. S1A, see time point 0), growth of cells containing the truncated allele was nearly abolished (Fig. 2A). In contrast, apparent residual expression of full-length *TOR1* was sufficient to permit substantial growth during *tetO* repression (Fig. 2A), possibly due to inevitable leakiness of an inhibited *tetO* construct. qRT-PCR showed rapid overexpression of *TOR1-Del381* alleles from *tetO* in the absence of doxycycline, while *TOR1-FL* mRNA levels recovered more slowly after *tetO* repression, hinting at a possible role of N-terminal HEAT repeats in the half-life of the *TOR1* mRNA itself (Fig. S1A). We assayed TORC1 activity during exponential growth in rich complex medium (YPD) by the phosphorylation state of ribosomal protein S6 (P-S6) [25], using a minimal doxycycline concentration of 5 ng/ml as a ceiling for the expression level. Cells overexpressing full-length *TOR1* or *TOR1-Del381* from *tetO* showed comparable P-S6 signal intensity to wild type cells in rich medium (Fig. 2B), with a slightly weaker P-S6 signal in Del381 cells under these conditions of optimal growth. Levels of total Rps6 protein did not change under any of the experimental conditions we examined and hence are shown only twice (Fig. 2B and D). We concluded that overexpression of *TOR1* from unrepressed *tetO* does not per se increase TORC1 signaling, possibly because the other TORC1 components required by the stoichiometry of the complex are not available at higher levels.

**Fig. 2.**
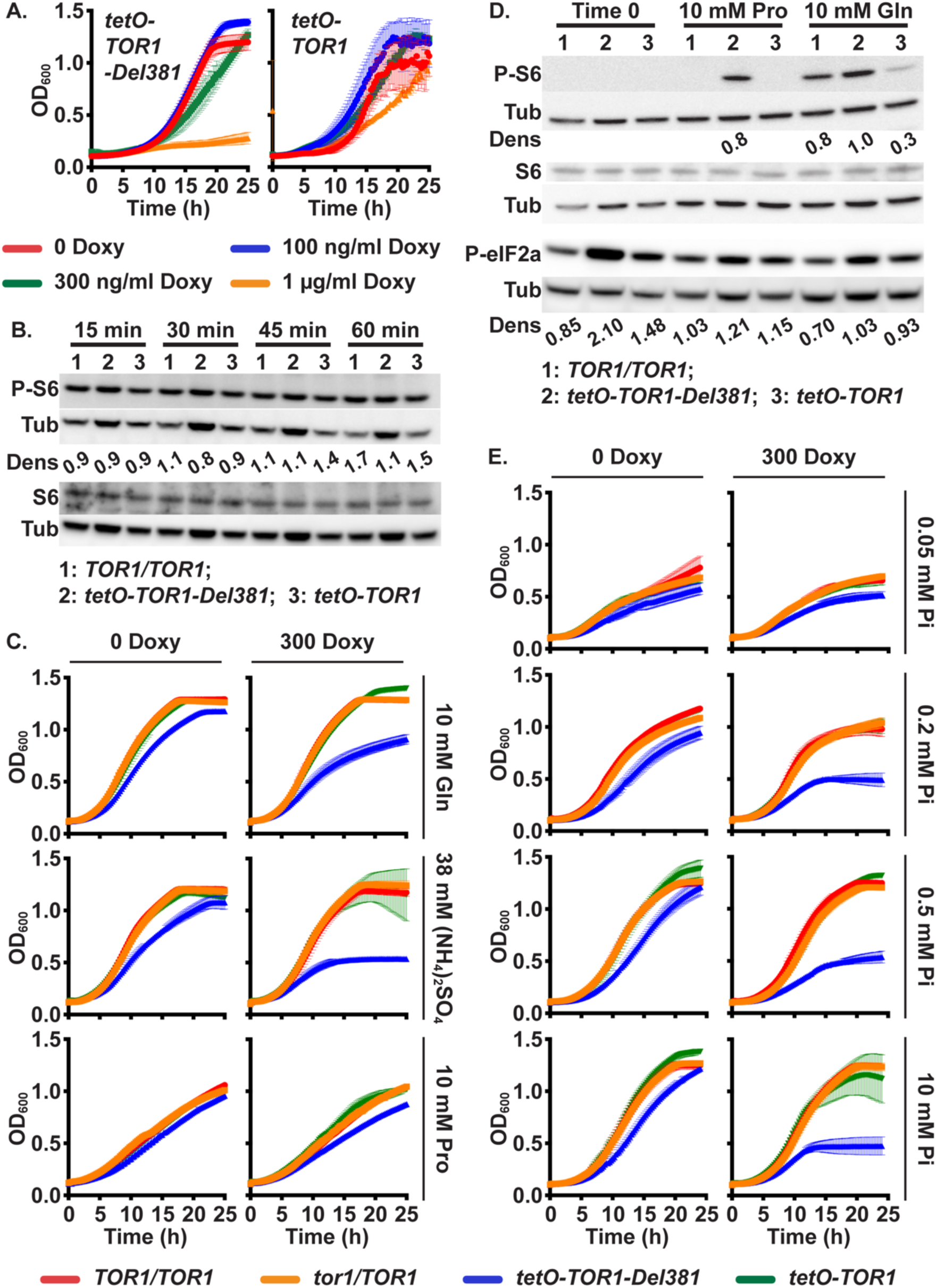
Tor1 N-terminal HEAT repeats are required to adapt growth velocity and TORC1 activity to availability of preferred nitrogen sources and of phosphate. **A.** Cells expressing *TOR1-Del381 or TOR1-FL* from *tetO* were pre-grown in YPD medium for 4 h, and inoculated in YPD with increasing concentrations of doxycycline (Doxy). OD_600_ was read every 15 minutes. **B.** Western blot of cells of indicated *TOR1* genotypes, wild type (*TOR1/TOR1*), Del381 and FL, grown in YPD with 5 ng/ml doxycycline; protein extracts probed with antibody to phosphorylated Rps6 (P-S6), total Rps6 (S6) and tubulin (Tub) as loading control. Dens: signal intensity ratio of P-S6 to Tub vs. signal intensity ratio of S6 to Tub. (*TOR1/TOR1*, JKC1713; *tetO-TOR1*-*Del381*, JKC1441; *tetO-TOR1,* JKC1549) **C.** Cells of indicated genotypes were grown in YNB without ammonium sulfate ((NH_4_)_2_SO_4_) supplemented with 10 mM Glutamine (Gln), 38 mM (NH_4_)_2_SO_4_ or 10 mM Proline (Pro) as sole nitrogen source, without or with 300 ng/ml doxycycline (300 Doxy). **D.** Western blot of cells pre-grown in YPD medium with 2 µg/ml doxycycline for 4 h, then incubated in YNB without doxycycline, without (NH_4_)_2_SO_4_, with 10 mM Proline (Pro) or Glutamine (Gln) as sole nitrogen source for 45 min; protein extracts probed with antibody to phosphorylated Rps6 (P-S6), total Rps6 (S6) or phosphorylated eukaryotic translation initiation factor 2A (P-eIF2*α*), and tubulin (Tub) as loading control. Dens: signal intensity ratio of P-S6 or P-eIF2*α* to Tub. Strains as in panel B and *TOR1/TOR1*, JKC1361. **E.** Cells pre-grown as in C were inoculated into YNB without phosphate supplemented with varying concentration of KH_2_PO_4_ (Pi), without or with 300 ng/ml doxycycline (300 Doxy). Error bars show SD of 3 technical replicates. Strains as in panel B and *tor1/TOR1*, JKC1347.

Preferred nitrogen sources activate *S. cerevisiae* TORC1 [43–45], and this effect is conserved in *C. albicans* [25]. To test the role of *C. albicans* Tor1 and its N-terminal HEAT repeats in cells’ responses to rich versus non-preferred nitrogen sources, wild type, *tor1/TOR1*, Del381 and FL cells were grown in different nitrogen sources. Without *tetO*-repression, or during repression in moderate concentrations of 300 ng/ml doxycycline, FL cells grew as well as wild type or *tor1/TOR1* heterozygous cells (Fig. 2C). In synthetic YNB medium with preferred nitrogen sources known to induce TORC1 signaling [25], ammonium sulfate or glutamine, Del381 strains grew more slowly than wild type or heterozygotes (Fig. 2C). In contrast, in the non-preferred nitrogen source proline [25], Del381 cells had no detectable specific growth defect while wild type and all *TOR1* genotypes grew more slowly than in preferred nitrogen sources (Fig. 2C). Cell dilutions of the same strains spotted on solid agar media with these different nitrogen sources, showed analogous phenotypes during overexpression of *tetO-TOR1-Del381* or *tetO-TOR1-FL* alleles in the absence of doxycycline, while during partial repression of these alleles with a moderate doxycycline concentration, growth of Del381 cells was sharply diminished on all 3 nitrogen sources (Fig. S1B). Tor1 N-terminal HEAT repeats were therefore specifically required for cells’ growth acceleration during their use of preferred nitrogen sources.

To examine the role of Tor1 N-terminal HEAT repeats in TORC1 signaling during growth in distinct nitrogen sources, we assayed TORC1 activity by the P-S6 signal [25]. Expression of *TOR1* was repressed for 4 hours by exposure to 2 µg/ml doxycycline and then released for 45 minutes in media without doxycycline containing glutamine or proline, respectively. Cell lysates were probed with antibody to P-S6 and total Rps6. Del381 cells showed significantly elevated P-S6 signals above those of wild type or FL cells in both conditions: in proline, where wild type and FL cells’ P-S6 signal was undetectable while that of Del381 cells was strong, and in glutamine, where the P-S6 signal from Del381 cells was more intense than that of the other two strains (Fig. 2D). Hence, downregulation of TORC1 signaling in response to a non-preferred nitrogen source required the complete Tor1 N-terminal HEAT domain.

Eukaryotic initiation factor 2 mediates translational control in response to starvation and environmental stresses in *S. cerevisiae* as reviewed e.g. in [46]. During starvation for preferred nitrogen sources and for glucose, the eIF2 alpha subunit (eIF2α) is phosphorylated by the kinase Gcn2 [47] and translation of many anabolic messenger RNAs is inhibited. Using an antibody against the conserved phospho-serine 51 of human eIF2α, we examined whether inappropriately increased TORC1 signaling in Del381 cells might correspond to inappropriately weak translation inhibition signaling through Gcn2, as assayed by eIF2α phosphorylation. To the contrary, we found that eIF2α phosphorylation was actually increased in Del381 cells (Fig. 2D), reflecting increased inhibitory signaling by Gcn2 and suggesting that TORC1- and Gcn2- signaling can become uncoupled when TORC1 lacks a function provided by N-terminal HEAT repeats.

### Tor1 N-terminal HEAT repeats were required for growth acceleration in phosphate-replete conditions

During invasion of the host, *C. albicans* apparently experiences starvation for inorganic phosphate, since expression of the high-affinity inorganic phosphate transporters Pho84 and Pho89 is induced in models of invasive disease [48–51], and since loss of Pho84 attenuates virulence [52]. We examined the role of Tor1 and its N-terminal HEAT repeats in distinct conditions of phosphate availability. Cells containing *TOR1* genotypes wild type, heterozygous null (*tor1/TOR1*), Del381 and FL alleles were grown in distinct phosphate concentrations. In low ambient phosphate, cells overexpressing *TOR1-Del381* in the absence of doxycycline grew at rates comparable to those overexpressing *TOR1-FL* (Fig. 2E); they showed mildly increasing growth defects with increases in phosphate concentrations (Fig. 2E). During moderate repression of *tetO*, the growth defect of Del381 cells became more pronounced in increasing ambient phosphate concentrations compared with cells containing full length *TOR1* alleles (Fig. 2E). We concluded that the N-terminal HEAT repeats were required for growth acceleration in favorable phosphate- as well as nitrogen source conditions.

The response of TORC1 signaling in wild type, Del381 and FL cells to external phosphate repletion or - starvation was examined. P-S6 was not decreased in low-phosphate (0.22 mM KH_2_PO_4_) versus phosphate-replete (11 mM KH_2_PO_4_) conditions in Del381 cells (Fig. S1C). Overall growth, as assayed by an increase in the optical density of the culture, was therefore delinked from the intensity of TORC1 signaling in Del381 cells.

### Tor1 N-terminal HEAT repeats contributed to acceleration of growth during glucose repletion and were required for growth on lactate and for physiologic oxygen consumption

Carbon source- and phosphate repletion are components of *C. albicans* cells’ nutrient status monitored by TORC1 [25, 33]. In the host interaction, *Candida* cells compete with host phagocytes for glucose [53–55]. During growth in low concentrations of glucose, cells overexpressing *TOR1-Del381* from *tetO* grew at similar rates as wild type or cells expressing full length *TOR1*. In high glucose concentrations, Tor1-Del381 expressing cells showed a slower growth rate than those containing an intact N-terminal HEAT domain (Fig. 3A) and this effect again became more pronounced during partial repression of *tetO* with a moderate doxycycline concentration of 300 ng/ml. This finding suggested that functions residing in the N-terminus of the N-terminal HEAT domain contributed to growth acceleration during repletion of the preferred carbon source, glucose.

**Fig. 3.**
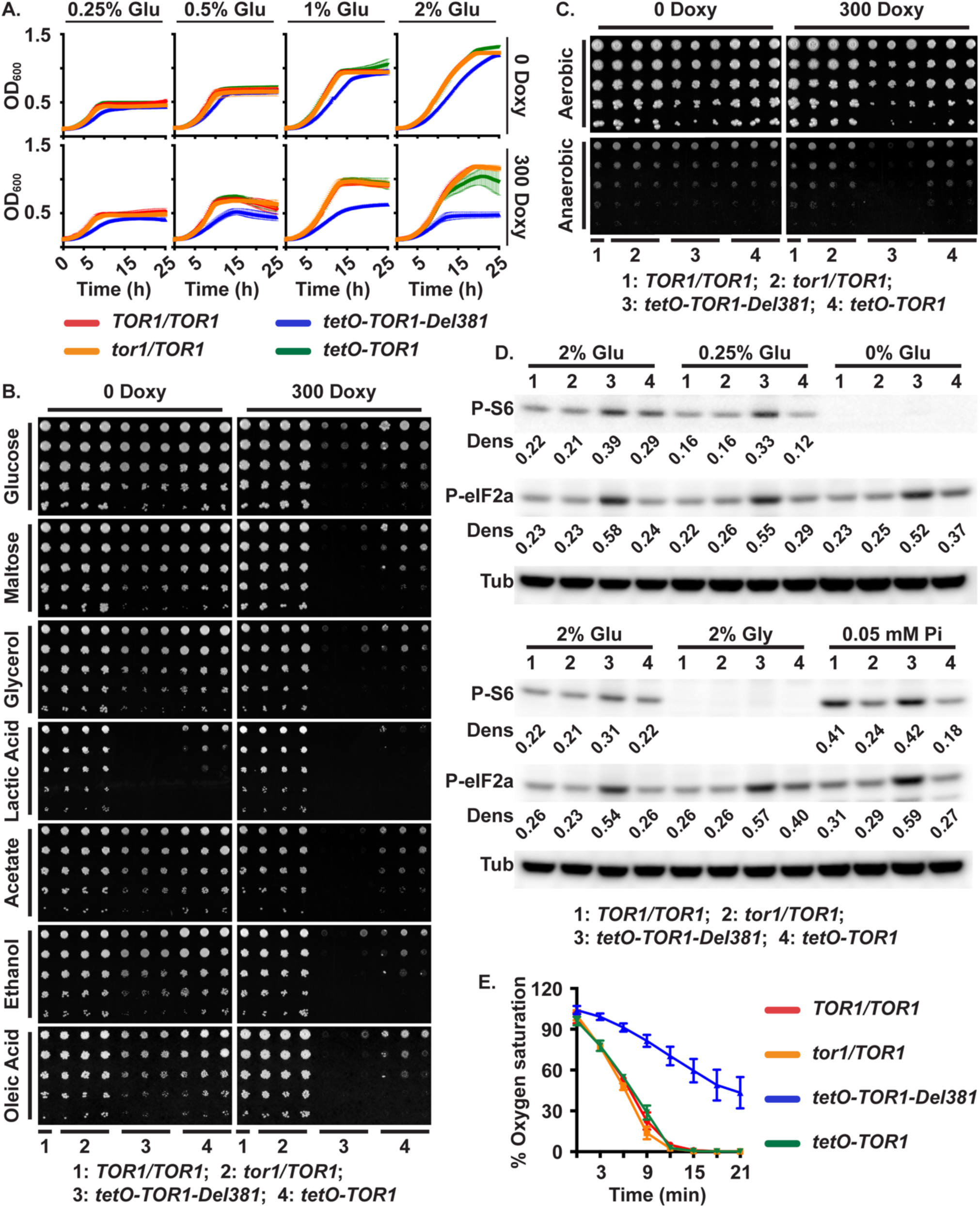
Cells lacking Tor1 N-terminal HEAT repeats fail to adapt growth to carbon source availability and are defective in growth on lactate and in oxygen consumption. **A.** Cells were grown in YNB medium with varying concentration of glucose (Glu), without or with 300 ng/ml doxycycline (300 Doxy). **B.** Cells of indicated genotypes were spotted onto YPD agar medium containing different carbon sources (2% w/v), without or with 300 ng/ml doxycycline. (*TOR1/TOR1*, JKC1713; *tor1/TOR1*, JKC1345, JKC1346, JKC1347; *tetO-TOR1*-*Del381*, JKC1442, JKC1445, JKC1441; *tetO-TOR1,* JKC1543, JKC1546, JKC1549). **C.** Cells of indicated genotypes were spotted onto YPD agar medium without or with 300 ng/ml doxycycline, and incubated either aerobically or anaerobically for 48 h. **D.** Western blot. Cells pre-grown in YPD with 5 ng/ml doxycycline for 4 h were inoculated into YNB without Pi with 5 ng/ml doxycycline containing nutrient concentrations equivalent to, or distinct from, standard YNB (2% Glu, 2% glucose, 10 mM Pi; 0.25% Glu, 0.25% glucose, 10 mM Pi; 0% Glu, no added direct carbon source, 10 mM Pi; 2% Gly, 2% glycerol, 10 mM Pi; 0.05 mM Pi, 2% glucose, 0.05 mM Pi). Protein extracts were probed with antibody to phosphorylated Rps6 (P-S6) or phosphorylated eukaryotic translation initiation factor 2A (P-eIF2*α*), and tubulin (Tub) as loading control. Dens: signal intensity ratio of P-S6 or P-eIF2*α* to Tub. (*TOR1/TOR1*, JKC1713; *tor1/TOR1*, JKC1346; *tetO-TOR1-Del381*, JKC1445; *tetO-TOR1,* JKC1546). **E.** Percentage of oxygen saturation of the medium in strains with distinct *TOR1* alleles grown in YPD. Error bars show SD of 3 biological replicates. (*TOR1/TOR1*, JKC1713; *tor1/TOR1*, JKC1347; *tetO-TOR1-Del381*, JKC1441; *tetO-TOR1,* JKC1549).

*C. albicans* does not encounter high glucose concentrations during invasive infection: human bloodstream glucose is 0.1%. *Candida*’*s* ability to use a range of carbon sources contributes to fitness in the host [53, 56–63]. The contribution of Tor1 and its N-terminal HEAT repeats to utilization of another fermentable carbon source, maltose, and of the non-fermentable carbon sources glycerol, ethanol, lactate, acetate, and oleic acid were examined. During full induction of *tetO* in the absence of doxycycline, Del381 cells had a significant growth defect on rich complex medium with lactate as the carbon source (Fig. 3B). On all other carbon sources their growth defects during *tetO* induction compared with FL cells were minor (Fig. 3B). During moderate repression of *tetO*, growth defects of Del381 cells were more severe than those expressing full length Tor1 (Fig. 3B). Del381 cells’ growth defect on lactate was not due to acid stress, because they grew equally well on glucose- or glycerol-containing medium buffered to pH 2, as on unbuffered rich medium (Fig. S2). Physiologic expression of full-length *TOR1* was required for appropriate carbon source use. N-terminal HEAT repeats were specifically required for growth on lactate.

Since utilization of fermentable carbon sources like glucose and maltose was not dramatically impaired in cells overexpressing *TOR1-Del381* from *tetO* (in the absence of doxycycline), compared with non-fermentable carbon sources like glycerol and ethanol (Fig. 3B), we asked whether these cells were able to grow anaerobically, i.e. under conditions requiring fermentation. Del381 cells again had no dramatic growth defect under anaerobic (hypoxic) conditions compared with wild type during overexpression of *TOR1-Del381* in the absence of doxycycline (Fig. 3C); only when *TOR1-Del381* transcription was moderately repressed were these cells unable to grow anaerobically (Fig. 3C). We concluded that fermentation does not require a regulatory activity residing in the Tor1 N-terminal HEAT repeats.

TORC1 activation and translational regulation through eIF2*α* were examined in cells growing in 2% glucose (control), 0.25% glucose, 2% glycerol and in the absence of a carbon source. The P-S6 signal, indicating TORC1 activation, was slightly reduced in the lower glucose concentration (Fig. 3D). Del381 cells showed aberrantly increased P-S6 intensity in 0.25% glucose, similarly to their behavior during nitrogen starvation. In the absence of a direct carbon source (0 glucose) and in 2% glycerol, the P-S6 signal was undetectable for all strains. Phosphorylation of eIF2*α* did not change during provision of different glucose concentrations or glycerol (Fig. 3D), but the P-eIF2*α* signal again was stronger in Del381 cells under these conditions, including when they had no detectable P-S6 signal. *C. albicans* Gcn2 signaling apparently did not respond to carbon source provision under these experimental conditions; it was uncoupled from TORC1 activation in cells lacking N-terminal HEAT repeats.

We asked whether lagging growth of Del381 cells in high glucose concentrations corresponded to decreased respiration. We measured oxygen consumption of wild type, *tor1/TOR1*, FL and Del381 cells. Oxygen consumption was significantly decreased in Del381 cells in which *tetO-TOR1-Del381* was not repressed, consistent with a slower respiratory metabolism (Fig. 3E). We concluded that N-terminal HEAT repeats contributed to increasing oxidative phosphorylation when glucose was abundant.

### Oxidative stress endurance required Tor1 N-terminal HEAT repeats

Reduced oxygen consumption of Del381 cells implies a lower rate of mitochondrial activity that generates reactive oxygen species (ROS). DCFDA-detectable ROS were in fact lower in cells with *tetO*-controlled *TOR1* alleles (Fig. 4A). We questioned whether decreased intracellular ROS concentrations of FL and Del381 cells might enable better endurance of oxidative stress; alternatively, perturbation of Tor1 could impair the switch from growth-promoting to stress-enduring processes and increase sensitivity to oxidative stress. We examined the role of Tor1 and its N-terminal HEAT repeats in the fungus’s endurance of oxidative stress by exposing cells to the superoxide-generating compound plumbagin and the peroxide-generating agent H_2_O_2_. *tor1/TOR1* heterozygous cells, spotted in 5-fold dilutions on YPD plates containing plumbagin or H_2_O_2_, were able to tolerate these compounds as well as wild type, but cells expressing *tetO-TOR1-FL* were hypersensitive to H_2_O_2_ in the absence and presence of *tetO* repression with doxycycline (Fig. 4B). Del381 cells were strikingly hypersensitive to both sources of ROS (Fig. 4B). These findings confirmed that TORC1 contributes to managing oxidative stress in *C. albicans*, and suggested that the N-terminal HEAT repeats of the Tor1 protein were critical for its role in oxidative stress endurance.

**Fig. 4.**
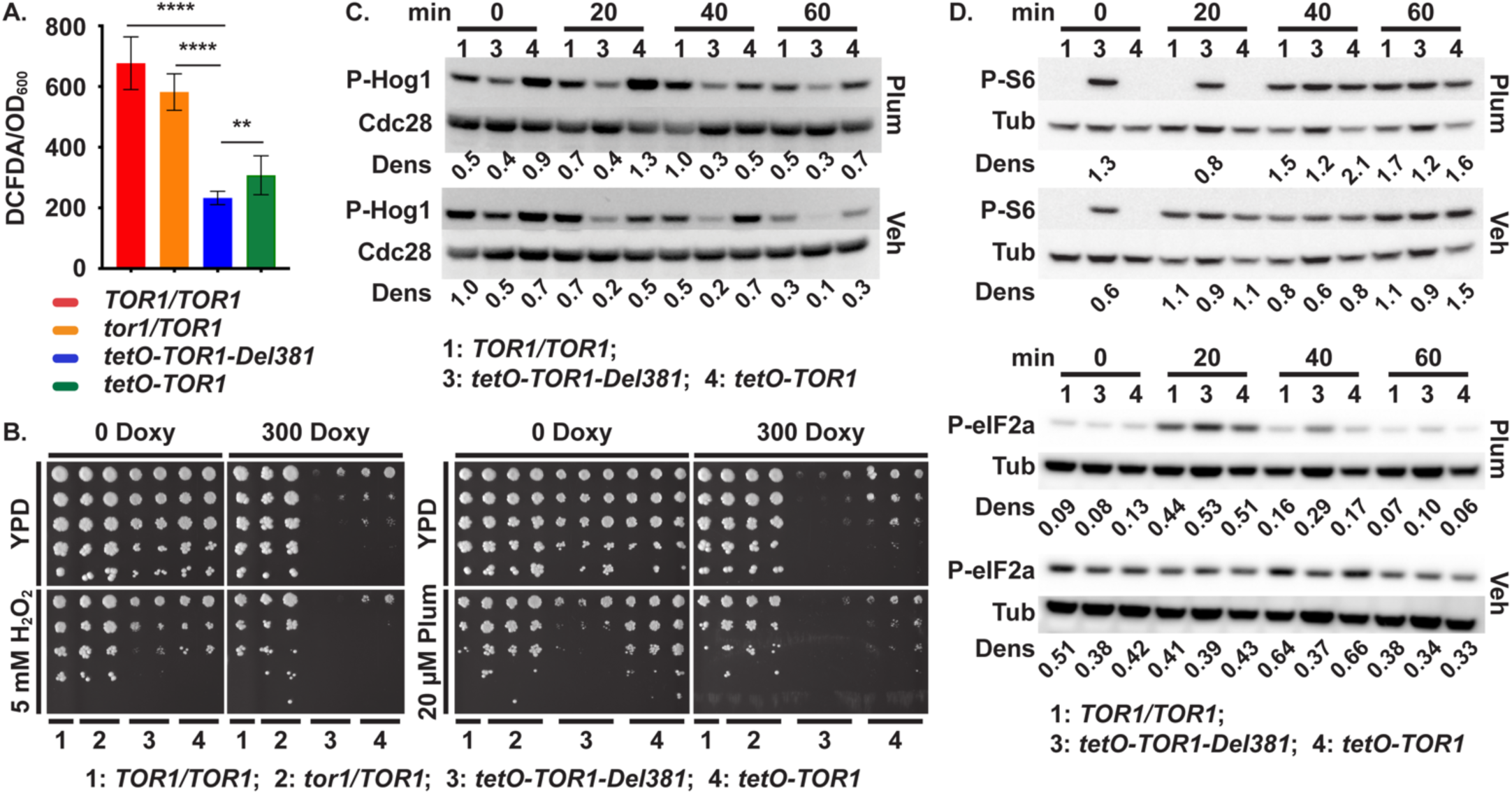
Tor1 N-terminal HEAT repeats are required for oxidative stress responses. **A.** DCFDA-detectable ROS. Cells cultured overnight in YPD were diluted in SC medium (LoFlo) at OD_600_ 0.5 and fluorescence intensity was determined after staining cells with 50 µM DCFDA for 90 minutes. **** is *p*<0.0001; ** is *p*=0.0079; error bars show SD of 3 biological replicates. (*TOR1/TOR1*, JKC1713; *tor1/TOR1*, JKC1347; *tetO-TOR1-Del381*, JKC1441; *tetO-TOR1,* JKC1549). **B.** Dilutions of cells of indicated genotypes were spotted on YPD medium with or without 300 ng/ml doxycycline (300 Doxy), oxidative stress was induced with 5 mM H_2_O_2_ or 20 µM Plumbagin (Plum). (*TOR1/TOR1*, JKC1361; *tor1/TOR1*, JKC1345, JKC1346, JKC1347; *tetO-TOR1-Del381*, JKC1442, JKC1445, JKC1441; *tetO-TOR1,* JKC1543, JKC1546, JKC1549). **C, D.** Cells of indicated genotypes were pre-grown in YPD medium with 5 ng/ml doxycycline for 3.5 h and diluted into fresh YPD medium with 5 ng/ml doxycycline with either 10 µM Plumbagin (Plum) or DMSO as vehicle (Veh). Total protein extract was probed with antibody to phosphorylated Hog1 (P-Hog1) and the PSTAIRE antigen of Cdc28 as loading control (**C**), or with antibody to phosphorylated Rps6 (P-S6) and eIF2*α* (P-eIF2*α*), and tubulin (Tub) as loading control (**D**). Dens: signal intensity ratio of P-Hog1 to Cdc28 (**C**) or P-S6 or P-eIF2*α* to Tub (**D**) (*TOR1/TOR1*, JKC1361 for **C** and JKC1713 for **D**; *tetO-TOR1-Del381*, JKC1441; *tetO-TOR1,* JKC1549).

Oxidative stress activates signaling through the HOG MAP kinase pathway in *C. albicans* as assayed by the Hog1 phosphorylation state [64–66]. This pathway induces antioxidant mechanisms like catalase, superoxide dismutase and enzymes involved in the thioredoxin and glutaredoxin systems [66]. Del381 cells were defective in activating Hog1 phosphorylation in response to plumbagin exposure, under conditions in which FL cells induced a strong phospho-Hog1 signal (Fig. 4C). This result suggested that Tor1 N-terminal HEAT repeats were required to induce a physiologic Hog1 oxidative stress response.

We tested whether during oxidative stress exposure, TORC1 signaling was physiologically downmodulated in these *tor1* mutants. Wild type and FL cells responded to plumbagin exposure as expected, by inhibiting Rps6 phosphorylation as evinced by an absent P-S6 signal on Western blot in the first 20 minutes of the time course. In contrast, Del381 cells failed to downmodulate P-S6, remaining in an abnormally activated TORC1 state even immediately after exposure to plumbagin (Fig. 4D). To exclude an effect of plumbagin on transcription from *tetO*, we examined *TOR1* mRNA levels by qRT-PCR in each of the strains and found no plumbagin effect (Fig. S3).

Since in *S. pombe*, translation initiation during oxidative stress is suppressed by Gcn2 kinase’s phosphorylation of eIF2α, we examined this response in *C. albicans* cells from the same cell lysates exposed to plumbagin or vehicle that were assayed for P-S6. In vehicle the P-S6 and P-eIF2α signals of Del381 cells were not substantially different from those of wild type or FL cells (Fig. 4D). In contrast, in plumbagin-treated cells, the P-eIF2α signal of Del381 cells was stronger than in wild type or FL cells, indicating stronger pro-inhibitory activity by Gcn2 (Fig. 4D). Hence in the absence of stress, Del381 cells’ TORC1- and Gcn2 signaling were aligned. During oxidative stress with plumbagin these pathways were uncoupled in these cells, with inappropriately increased TORC1 signaling and increased counter-regulation by inhibitory Gcn2 signaling.

### Downregulation of translation during oxidative stress required Tor1 N-terminal HEAT repeats

Translation initiation of most messages [29] is induced during TORC1 activation and repressed during TORC1 inhibition, with specific regulatory mechanisms known for mTOR [67]. During oxidative stress, as during TORC1 inhibition by rapamycin, translation initiation in *S. cerevisiae* is also inhibited through Gcn2-dependent phosphorylation of eIF2α, as well as through decreased ribosomal transit [68]; Gcn2 also responds to oxidative stress to inhibit translation initiation in *S. pombe* [69]. Since TORC1 signaling as reflected in the P-S6 signal was hyperactive in Del381 cells during oxidative stress, while paradoxically, their translation inhibition through Gcn2 phosphorylation of eIF2*α* was also increased, we questioned which signaling activity determined the final output, translational activity. We used a heterologous message, GFP, whose transcription we could control from *tetO* and which presumably had no internal sequences directing translational regulation. We examined whether appearance of the protein was decreased in *C. albicans* during exposure to the superoxide generating compound plumbagin (Fig. 5A). As expected, wild type cells expressing GFP from inducible *tetO* showed slower appearance of a GFP signal during exposure to plumbagin than to vehicle (Fig. 5A).

**Fig. 5.**
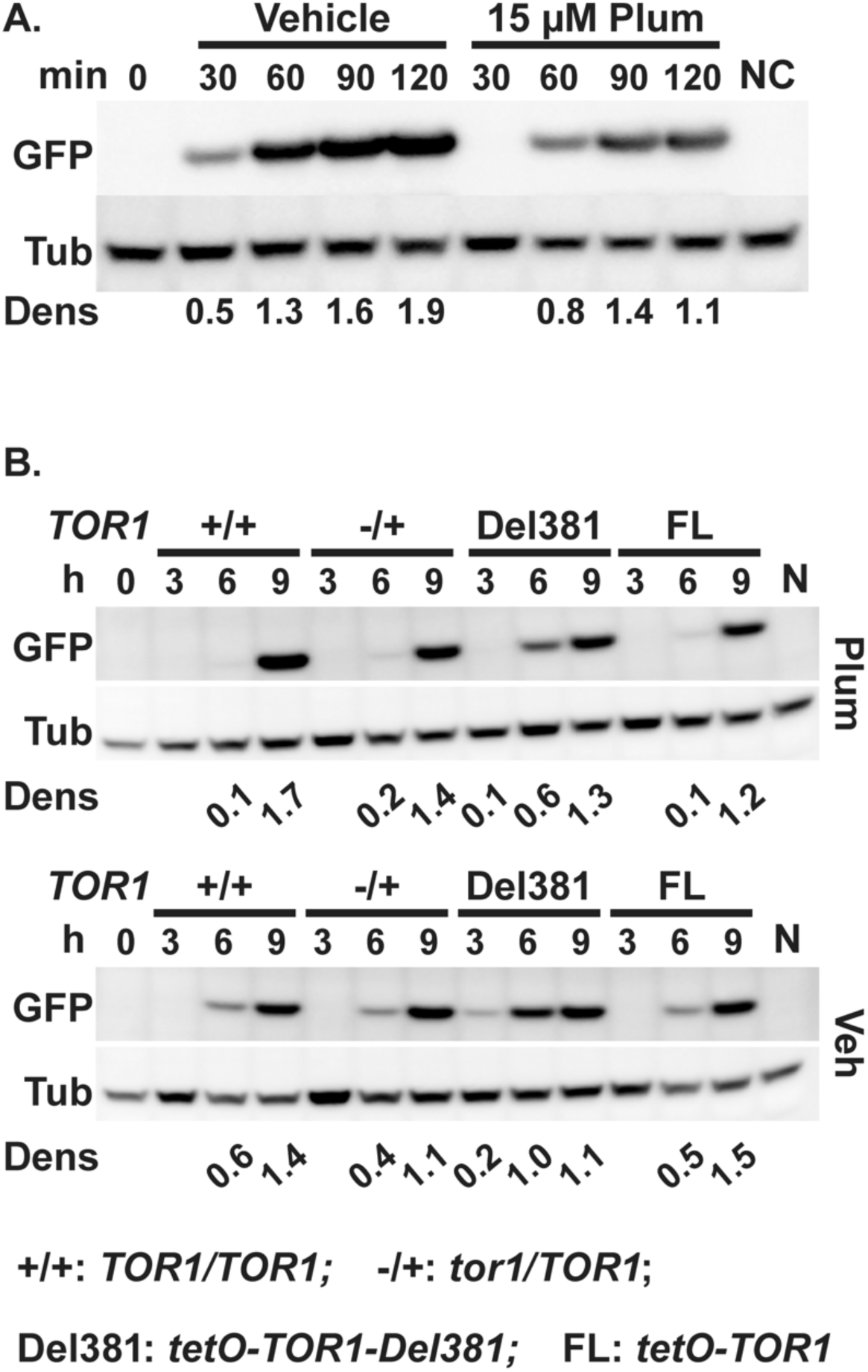
Tor1 N-terminal HEAT repeats are required for oxidative stress-induced delay of translation initiation. **A.** Cells expressing GFP from *tetO* (ON) (strain TETG25B) were grown in YPD medium for 15 h, then inoculated into fresh YPD and pre-grown for 3.5 h (Time 0, 0 min). GFP expression was induced with 50 µg/ml doxycycline in SC medium with Vehicle (DMSO) or 15 µM Plumbagin (Plum). Negative control (NC) cells were grown in SC medium without doxycycline (with DMSO) for 120 min. Total protein extracts were probed with antibody to GFP, and tubulin (Tub) as loading control. Dens: signal intensity ratio of GFP to Tub. Representative of 2 biological replicates. **B.** Cells expressing *pMAL2*-GFP in backgrounds with distinct *TOR1* alleles were pre-grown in YPD medium with 10 ng/ml doxycycline for 15 h (+/+, JKC2616; -/+, JKC2620; Del381, JKC2624; FL, JKC2628). GFP expression was induced by inoculation into YP-Maltose medium with 5 ng/ml doxycycline, containing 13 µM Plumbagin (Plum) or DMSO as vehicle (Veh). (0 h, JKC2616 after pre-growth; N, JKC2616 inoculated into YPD instead of YP-Maltose with 5 ng/ml doxycycline, Plumbagin or DMSO, grown for 9 h). Total protein extracts were probed with antibody to GFP, and tubulin (Tub) as loading control. Dens: signal intensity ratio of GFP to Tub.

The effect of plumbagin on GFP translation in cells containing distinct *TOR1* alleles in which GFP was expressed from the conditional *MAL2* promoter (*pMAL2*) was then examined. When cells are shifted from glucose to maltose, the *MAL2* promoter is induced; we assayed appearance of a GFP protein signal after this shift in cells containing *TOR1* genotypes wild type, *tor1/TOR1,* Del381 and FL. Translation of GFP to detectable levels occurred earlier in plumbagin- and vehicle-exposed Del381 cells than in FL cells (Fig. 5B). This finding indicated that translation was aberrantly upregulated in Del381 cells and that the hyperactive TORC1 signal, not simultaneously increased eIF2*α* phosphorylation and hence inhibition by Gcn2, determined this final output.

### Tor1 N-terminal HEAT repeats participated in the response to cell wall- and heat stress

Host phagocytes exert physical force and break *C. albicans* hyphal cells [70]. Mechanical stress and weakening of cell walls induces protein kinase C (PKC)-dependent cell wall integrity responses to prevent *C. albicans* cells from rupturing or breaking [71–74]. Activation of these responses can be assayed by the Mkc1 phosphorylation state [73]. In *S. cerevisiae*, an upstream cell wall integrity pathway component Rho1, in its GTP-loaded state, activates the major TORC1 effector Tap42-2A phosphatase that promotes stress responses by displacing it from its binding site on Kog1 [75]. In a modulatory loop, Tor kinase function is required for *S. cerevisiae* Rho1 activity, while inhibition of TORC1 by rapamycin induces cell wall integrity signaling [75]. We examined the ability of cells lacking Tor1 N-terminal HEAT repeats to respond to cell wall stress. Low concentrations of the beta-1,3-glucan synthase inhibitor micafungin, which induces cell wall stress, were strongly inhibitory to these cells on agar medium even when *tetO-TOR1-Del381* expression was induced in the absence of doxycycline (Fig. 6A). The cell wall disrupting dye Congo red, which binds to chitin fibrils [76], similarly had a very strong inhibitory effect on Del381 cells on agar media, regardless of induction or inhibition of their *tetO-TOR1-Del381* allele (Fig. 6A). Reexamining this phenotype in liquid media, given the activation of cell wall integrity pathway signaling by cells’ contact with agar surfaces [73], we found that *TOR1-Del381-*expressing cells were extremely sensitive to micafungin, whether the allele was overexpressed in 0 doxycycline or moderately repressed in 300 ng/ml doxycycline; *TOR1-FL-*expressing cells were hypersensitive during *tetO* repression but to a lesser extent (Fig. 6B). To confirm that this strong phenotype was not specific to the drug or the cell wall component inhibited (beta-1,3-glucan), we exposed the cells to the chitin synthase inhibitor nikkomycin. Since nikkomycin competes with components of YPD for plasma membrane uptake through oligopeptide transporters, we used synthetic complete medium (SC) for these experiments. Del381 and, to a lesser extent, FL cells were also hypersensitive to nikkomycin during moderate *tetO* repression, though the difference to wild type cells’ inhibition was less stark (Fig. 6B). The Tor1 N-terminal HEAT domain hence had a distinct role in responding to cell wall stressors.

**Fig. 6.**
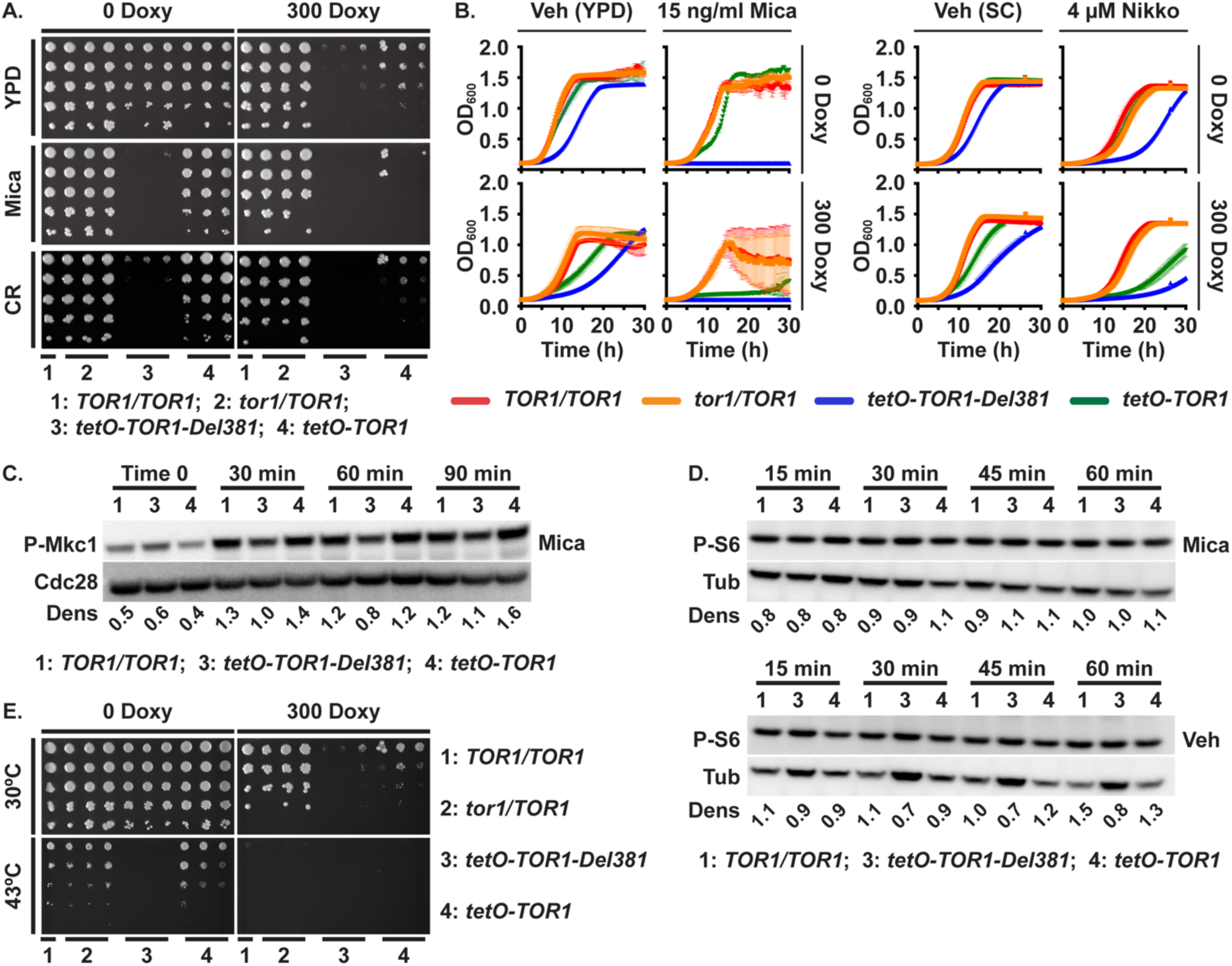
Tor1 N-terminal HEAT repeats are required for adequate cell wall- and heat stress responses. **A, E.** Dilutions of cells of indicated genotypes were spotted on YPD medium without or with 300 ng/ml doxycycline (300 Doxy), cell wall stress was induced with 10 ng/ml micafungin (Mica) or 15 µg/ml Congo red (CR) (**A**), and heat stress was induced at 43° (**E**); strains *TOR1/TOR1*, JKC1361 for YPD and micafungin and JKC1713 for Congo red and heat stress; *tor1/TOR1*, JKC1345, JKC1346, JKC1347; *tetO-TOR1-Del381*, JKC1442, JKC1445, JKC1441; *tetO-TOR1,* JKC1543, JKC1546, JKC1549). **B.** Cells of indicated genotypes were grown in YPD medium containing Vehicle (H_2_O) or 15 ng/ml micafungin (Mica), without or with 300 ng/ml doxycycline (left panel); or in synthetic complete medium (SC) containing Vehicle (H_2_O) or 4 µM nikkomycin (Nikko), without or with 300 ng/ml doxycycline (right panel). Strains *TOR1/TOR1*, JKC1713; *tor1/TOR1*, JKC1347; *tetO-TOR1-Del381*, JKC1441; *tetO-TOR1,* JKC1549. Error bars show SD of 3 technical replicates. **C, D.** Cells of indicated genotypes were pre-grown in YPD medium with 5 ng/ml doxycycline for 3.5 h (Time 0) and diluted into fresh YPD medium with 5 ng/ml doxycycline with either 10 ng/ml micafungin (Mica) or H_2_O as vehicle (Veh, **D** lower panel is the same P-S6 blot as Fig. 1B). Total protein extract was probed with antibody to phosphorylated Mkc1 (P-Mkc1) and the PSTAIRE antigen of Cdc28 as loading control (**C**), or with antibody to phosphorylated Rps6 (P-S6) and tubulin (Tub) as loading control (**D**). Dens: signal intensity ratio of P-Mkc1 to Cdc28 (**C**) or P-S6 to Tub (**D**); strains *TOR1/TOR1*, JKC1713; *tetO-TOR1-Del381*, JKC1441; *tetO-TOR,* JKC1549. Representative of 2 biological replicates.

Cell wall integrity signaling can be assayed in *C. albicans* by the Mkc1 phosphorylation state [73]. The P-Mkc1 signal intensity was minimally weaker in Del381 than in FL cells during micafungin exposure, though not at baseline (Figs. 6C, S4A), suggesting that the strong growth defect of these cells during cell wall stress was not related to a specific role of the Tor1 N-terminal HEAT domain in activating this pathway. Micafungin exposure did not affect Rps-6 phosphorylation even in wild type cells (Fig. 6D), confirming our previous observation that TORC1 downmodulation is not part of the physiologic response to cell wall stress [25]. Del381 cells’ hypersensitivity to cell wall stress therefore was not directly related to defective cell wall integrity responses.

Heat stress is encountered by *C. albicans* during invasive disease as a component of the host immune response [77]. In *S. cerevisiae*, GTP-loaded Rho1 promotes heat stress resistance by displacing Tap42-2A phosphatase from its inactive, Kog1-bound state, thereby releasing it into the cytoplasm and activating it [75]. However, it is the Kog1 N-terminus that binds Tap42 [75], while the Tor1 N-terminal HEAT domain is thought to interact with the Kog1 C-terminus in TORC1 [16]. To test whether Tor1 N-terminal HEAT repeats participate in the response to heat stress, we grew cells at 43° C in the absence and presence of sorbitol osmotic rescue which can separate direct heat stress from cell wall stress caused by heat. Del381 cells failed to grow at this elevated temperature, regardless whether *tetO* was induced or repressed (Fig. 6E). Growth defects at 43°, most severe in Del381 cells, were not osmotically rescued by 1 M sorbitol in any genotype (Fig. S4B), indicating heat stress hypersensitivity in cells with dysregulated Tor1. Hence a signal conveyed through Tor1 N-terminal HEAT repeats is required in the heat stress response.

During a 60-minute exposure to 41°, neither wild type nor *tor1* mutant cells downregulated Rps6 phosphorylation (Fig. S4C), indicating that this translation-inhibitory function is not part of the heat stress response. This finding also suggests that it was not lack of translation inhibition that left Del381 cells unable to tolerate elevated temperatures at which cells expressing a wild type *TOR1* allele could grow.

### Lack of N-terminal HEAT repeats sensitized cells to rapamycin and delayed their PKC-dependent rapamycin stress responses

Given Del381 cells’ inappropriate upregulation of translation during stress, we examined their response to direct inhibition of the Tor1 catalytic site with rapamycin. These cells were hypersensitive to rapamycin on solid and in liquid medium compared with cells carrying all other *TOR1* alleles, whether or not *tetO* was repressed (Fig. 7A,B, S4D). Rps6 phosphorylation was weaker during rapamycin exposure in Del381 cells at every time point we examined (Fig. 7C), indicating rapamycin hypersensitivity of anabolic TORC1 signaling in these cells, possibly due to distortion of the FRB site at which the complex of rapamycin and FKBP-12 docks onto Tor1 to obstruct substrate access to the catalytic cleft.

**Fig. 7.**
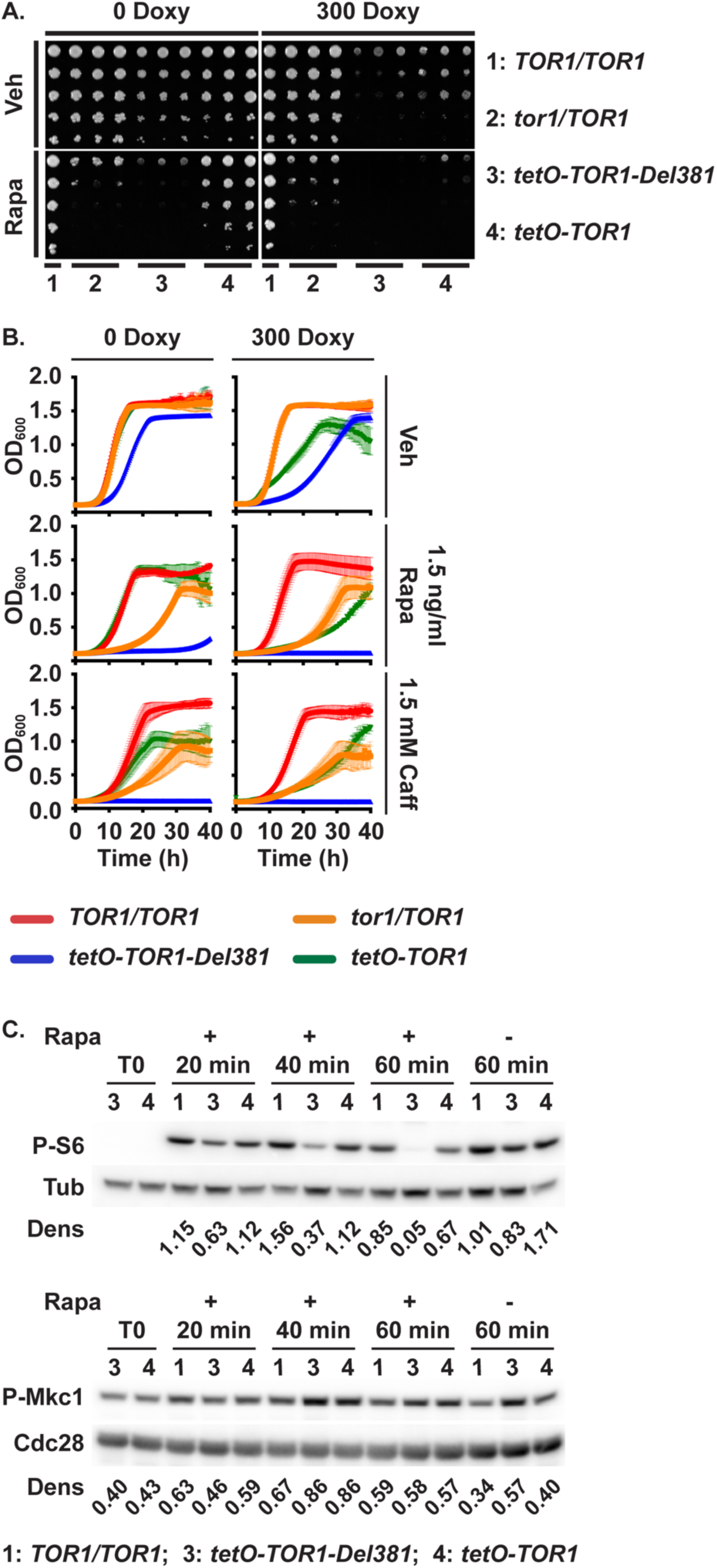
Cells lacking Tor1 N-terminal HEAT repeats are rapamycin hypersensitive while their cell wall integrity responses to rapamycin are delayed. **A.** Dilutions of cells of indicated genotypes were spotted on YPD medium containing vehicle (Veh, 90% ethanol) or 1.5 ng/ml rapamycin (Rapa), without or with 300 ng/ml doxycycline and incubated at 30° for 2 days. (*TOR1/TOR1*, JKC1713; *tor1/TOR1*, JKC1345, JKC1346, JKC1347; *tetO-TOR1-Del381*, JKC1442, JKC1445, JKC1441; *tetO-TOR1*, JKC1543, JKC1546, JKC1549) **B.** Cells of indicated genotypes were grown in YPD medium containing Vehicle (Veh, 90% ethanol), 1.5 ng/ml Rapamycin (Rapa) or 1.5 mM Caffeine (Caff), without or with 300 ng/ml doxycycline. Error bars show SD of 3 technical replicates. (*TOR1/TOR1*, JKC1713; *tor1/TOR1*, JKC1347; *tetO-TOR1-Del381*, JKC1441; *tetO-TOR1*, JKC1549). **C.** Cells of indicated genotypes were pre-grown in YPD medium with 5 ng/ml doxycycline for 3.5 h (Time 0, T0) and diluted into fresh YPD medium with 5 ng/ml doxycycline and 25 ng/ml rapamycin. Protein extracts were probed with antibody to phosphorylated Mkc1 (P-Mkc1) and the PSTAIRE antigen of Cdc28 as loading control, or with antibody to phosphorylated Rps6 (P-S6) and tubulin (Tub) as loading control. Dens: signal intensity ratio of P-Mkc1 to Cdc28 or P-S6 to Tub. *TOR1/TOR1*, JKC1713; *tetO-TOR1-Del381*, JKC1441; *tetO-TOR1*, JKC1549.

Caffeine inhibits Tor1 in a manner that is distinct from rapamycin’s mechanism of action but that appears to also involve the FRB domain [78]; in vitro caffeine inhibits ScTor1 kinase activity at an IC50 of 0.28 mm. We examined responses of Del381 and FL cells to caffeine exposure, in comparison with wild type and *tor1/TOR1* heterozygous null mutant cells. In contract to cells bearing the other 3 *TOR1* genotypes, Del381 cells were severely hypersensitive to caffeine whether their *TOR1-Del381* allele was overexpressed or repressed from *tetO* (Fig. 7B), consistent with the idea that Tor1 FRB domain function is affected when the N-terminal HEAT repeats are absent, though other mechanisms are possible.

Rapamycin exposure induces the PKC pathway stress response that can be assayed by Mkc1 phosphorylation intensity in *C. albicans* [25]. Del381 cells responded more slowly to rapamycin with increased P-Mkc1 signal intensity, and the increased signal receded more slowly, than in wild type and FL cells (Fig. 7C), suggesting that N-terminal HEAT repeats may be involved in down- as well as upregulation of the PKC pathway stress response during rapamycin exposure.

### Plasma membrane stress induced translation inhibition but Tor1 HEAT repeats were not required for membrane stress endurance

*C. albicans* is exposed to cytoplasmic membrane stress in the host e.g. by bile [79, 80] or by antimicrobial peptides [81]. We used low concentrations of SDS (0.005-0.01%) to test whether Tor1 and its N-terminal HEAT repeats are required in the response to membrane stress. During induction of *tetO-TOR1-Del381* in the absence of doxycycline, Del381 cells grew apparently normally on agar medium containing 0.005% SDS; only during repression of *tetO* by doxycycline did residual low-level *TOR1-Del381* expression fail to support growth on this medium (Fig. 8A).

**Fig. 8.**
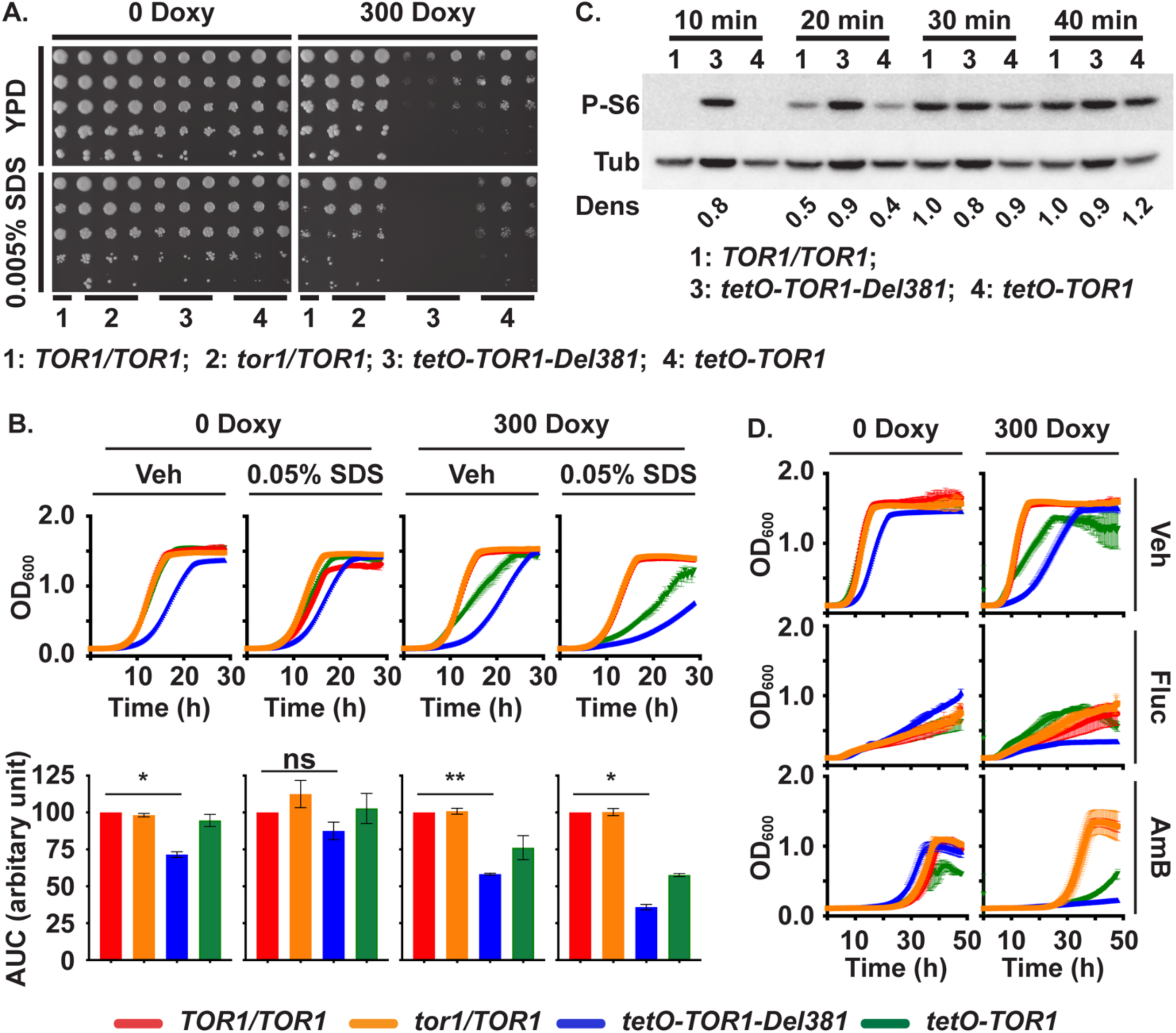
Plasma membrane stress responses do not significantly depend on Tor1 N-terminal HEAT repeats. **A.** Dilutions of cells of indicated genotypes were spotted on YPD medium without or with doxycycline (300 Doxy) and plasma membrane stress was induced with 0.005% SDS. (*TOR1/TOR1*, JKC1713; *tor1/TOR1*, JKC1345, JKC1346, JKC1347; *tetO-TOR1-Del381*, JKC1442, JKC1445, JKC1441; *tetO-TOR1,* JKC1543, JKC1546, JKC1549). **B.** Cells of indicated genotypes were grown in YPD medium containing Vehicle (Veh, H_2_O) or 0.05% SDS, without or with 300 ng/ml doxycycline. Upper panel shows the actual growth curves and lower panel shows the corresponding area under the curve (AUC) for each strain. ** is *p*=0.0076; * is *p*=0.029 (Veh, 0 Doxy) and *p*=0.0129 (0.05% SDS, 300 Doxy); ns is *p*=0.2, error bars show SD of 2 biological replicates (*TOR1/TOR1*, JKC1713; *tor1/TOR1*, JKC1347; *tetO-TOR1-Del381*, JKC1441; *tetO-TOR1,* JKC1549). **C.** Cells of indicated genotypes were pre-grown in YPD medium with 5 ng/ml doxycycline for 3.5 h and diluted into fresh YPD medium with 5 ng/ml doxycycline and 0.01% SDS. Total protein extract was probed with antibody to phosphorylated Rps6 (P-S6) and tubulin (Tub) as loading control. Dens: signal intensity ratio of P-S6 to Tub; strains as in panel B. **D.** Cells were grown as in B but treated with 1 μg/ml Fluconazole (Fluc) or 0.2 μg/ml Amphotericin B (AmB).

Even more striking than on solid media, cells overexpressing *tetO-TOR1-Del381* had a smaller growth defect, compared with wild type, in liquid SDS-containing medium than in vehicle (Fig. 8B), i.e. their relative growth defect was partially rescued by SDS exposure. Membrane stress exposure by 0.01% SDS in liquid medium induced downregulation of Rps6 phosphorylation in wild type and FL cells while Del381 cells were defective in this response (Fig. 8C). Repression of translation hence appeared to be a non-critical component of the physiologic response to plasma membrane stress, since Del381 cells’ defect in this response did not correspond to a growth defect.

Antifungal drugs like fluconazole and amphotericin B perturb plasma membrane function by inhibiting ergosterol biosynthesis or binding ergosterol. Cells overexpressing *tetO-TOR1-Del381* in the absence of doxycycline were not hypersensitive to these agents (Fig. 8D); their growth defect compared to wild type cells may, if anything, have been less pronounced during exposure to fluconazole and amphotericin B than during growth in vehicle (Fig. 8D), reminiscent of their growth in SDS. Taken together, these findings indicated that the role of N-terminal HEAT repeats in plasma membrane stress endurance was minor and separable from their role in controlling translation.

### Perturbation of Tor1 expression and loss of its N-terminal HEAT repeats led to hyphal growth misregulation with distinct phenotypes dependent on environmental conditions

Hyphal growth and -morphology is a result of multiple signaling pathways [82–84]. TORC1 signaling is known to affect *C. albicans* morphogenesis [37–40]. We examined hyphal growth on several filamentation-inducing media including M199 (Fig. 9A), Spider and RPMI (not shown); mutants in *TOR1* invariably had defective filamentation. Del381 and FL cells had decreased hyphal growth during overexpression of their *TOR1* alleles in 0 doxycycline, and during their moderate repression (Fig. 9A). On M199 medium of low pH during anaerobic growth, the hyphal growth defect of Del381 > FL cells was partially rescued compared to neutral pH (Fig. 9A). During anaerobic growth, cells overexpressing *TOR1-Del381* and *TOR1-FL* from *tetO* were hyperfilamentous, while moderate repression of *tetO* resulted in a hyperfilamentous phenotype of *TOR1-FL* expressing cells (Fig. 9A). Together these results indicated that the Tor1 N-terminal HEAT repeats, like the entire Tor1 protein, participated in a variety of signaling events whose final output is hyphal growth and which are modulated depending on multiple external and internal cellular conditions.

**Fig. 9.**
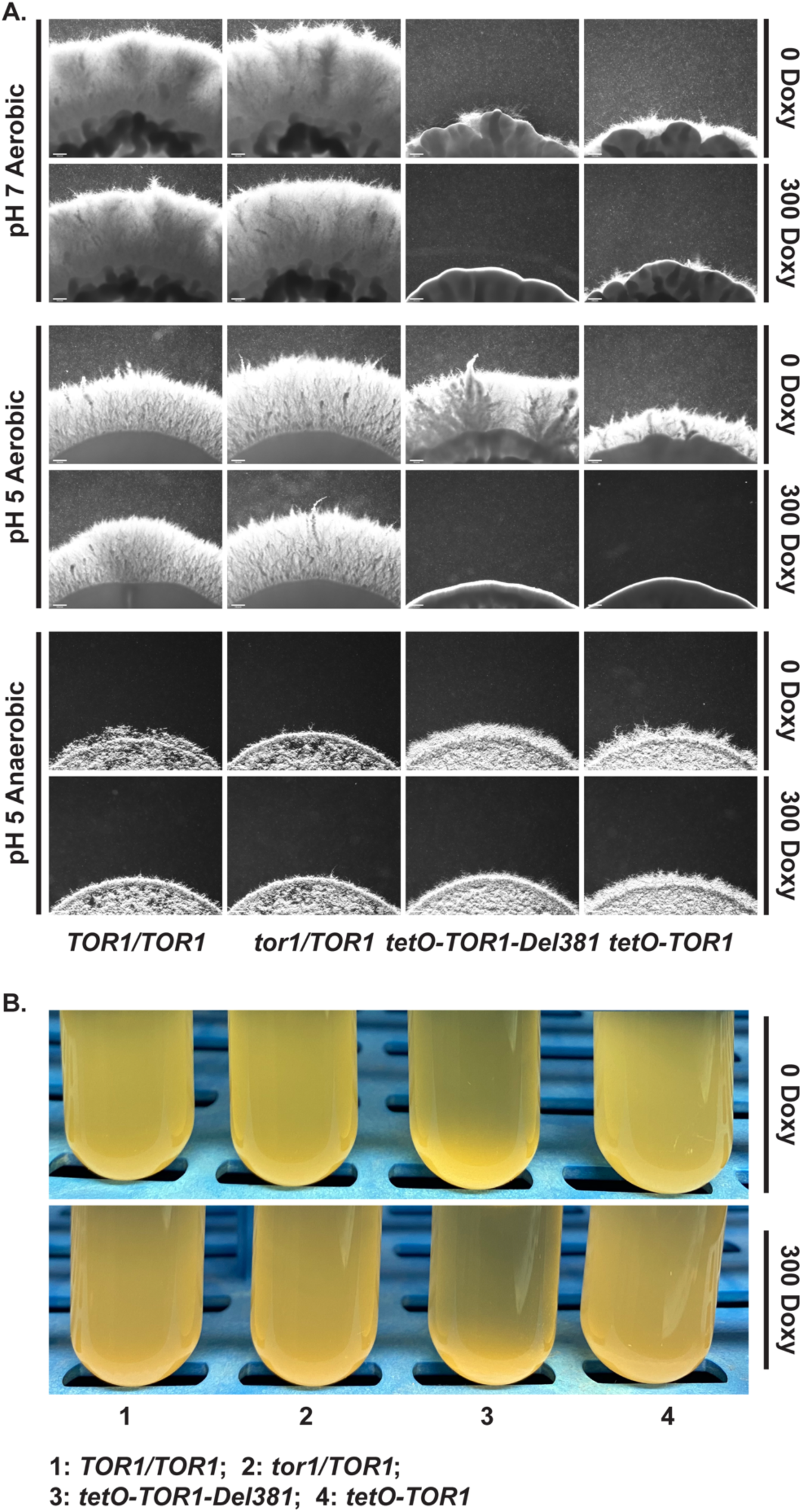
Filamentation responses to distinct stimuli depend on Tor1 N-terminal HEAT repeats. **A.** Cells of indicated genotypes were spotted at equidistant points around agar media, M199 pH 7 165 mM MOPS; M199 pH 5 100 mM MES, and incubated at 37° in aerobic or anaerobic conditions, without and with 300 ng/ml Doxycycline (300 Doxy). Spot edges were imaged. Scale bar 200 µm. **B.** Cells of indicated genotypes were pre-grown in YPD medium with 100 ng/ml doxycycline until exponential phase for 4 h. Flocculation was induced in Spider medium without or with 300 ng/ml Doxycycline (*TOR1/TOR1*, JKC1713; *tor1/TOR1*, JKC1347; *tetO-TOR1-Del381*, JKC1441; *tetO-TOR1,* JKC1549).

TORC1 inhibition with rapamycin was previously found to induce aggregation of *C. albicans* cells in liquid Spider medium through induction of adhesin gene expression [38]. We questioned whether Tor1 N-terminal HEAT repeats play a role in aggregation. Del381 cells aggregated excessively even during derepression of *tetO*-*TOR1-Del381* in the absence of doxycycline (Fig. 9B); aggregation was somewhat more pronounced during moderate *tetO* inhibition with 300 ng/ml doxycycline (Fig. 9B). Hence Tor1 N-terminal HEAT repeats contribute to repressing aggregation during physiological TORC1 activity.

Colony surface wrinkles are a filamentation phenotype characterized by Homann et al. for transcriptional regulator mutants [85]. On YPD+10% serum agar medium and on Spider medium, cells containing each of the *TOR1* alleles, in strains from different heterozygous lineages, had reproducible surface wrinkling phenotypes, each of which was distinctive for the mutant allele and for the agar medium (Fig. S5). We concluded that Tor1 N-terminal HEAT repeats had an important role in signaling events that control filamentation phenotypes.

### Cells lacking N-terminal HEAT repeats showed deregulation of multiple transcriptional modules comprising metabolism, stress responses and the white cell type

Microarray analysis was used to examine the transcriptome of Del381 cells compared to heterozygous *tor1/TOR1* cells in the absence of doxycycline repression. Gene Set Enrichment Analysis (GSEA) as described by Uwamahoro et al. [86] and Sellam et al. [87] showed that Del381 cells exhibited increased expression of genes involved in Ribosome Biogenesis and Assembly (GO:0042254) and Translation (GO:0006412), in line with their inappropriately upregulated Rps6 phosphorylation and GFP translation phenotypes. Deregulated expression of some genes previously associated with stress responses, including oxidative stress [66] and the response to hypoxia [87] was also observed (Fig. 10A, Supplementary Files 1 and 2). Genes controlled by the regulators of glucose- and carbohydrate metabolism Tye7 and Gal4 exhibited decreased expression [88]. While ribosome biogenesis genes were increased (*NOP4* and *6*, *RRP8, DIP2*, C1_04710C), paradoxically, starvation responses were also increased, such as components of the glyoxylate cycle (*ICL1, MLS1*), urea degradation (*DUR1* and *3*) and arginine degradation (*CAR1* and *CAR2*) pathways, import of glucose (*HGT1, 2, 8, 14* and *16*), of amino acids and oligopeptides (*CAN1* and *2, OPT1* and *2*) and of nucleobases and their precursors (*FUR4, XUT1*) with a concomitant decrease in amino acid biosynthetic genes like *MET15* and *16*, *ARG3*, *LYS1*, *9* and*12*, *HIS4*, *5* and *7* (Supplementary Files 1 and 2). Similar patterns were seen when examining genes whose expression increased during repression of *tetO-TOR1-Del381* in doxycycline, again reflecting genes whose physiologic repression was abnormally released in the absence of N-terminal HEAT repeats (Supplementary File 3). The genes whose expression level decreased during repression of *TOR1-Del381* in doxycycline had far lower amplitudes of fold-change than those whose expression increased (Supplementary Files 3 and 4), suggesting transcriptional downmodulation as a major role of regulatory outputs from the first 8 N-terminal HEAT repeats. Indicating a role for TORC1 and specifically N-terminal HEAT repeats in the white-opaque switch, *WH11* was among the most highly upregulated genes in Del381 cells compared with *tor1/TOR1* cells (Supplementary File 1), and during repression of both *TOR1-Del381* and of *TOR1-FL* with doxycycline (Supplementary Files 3 and 5).

**Fig. 10.**
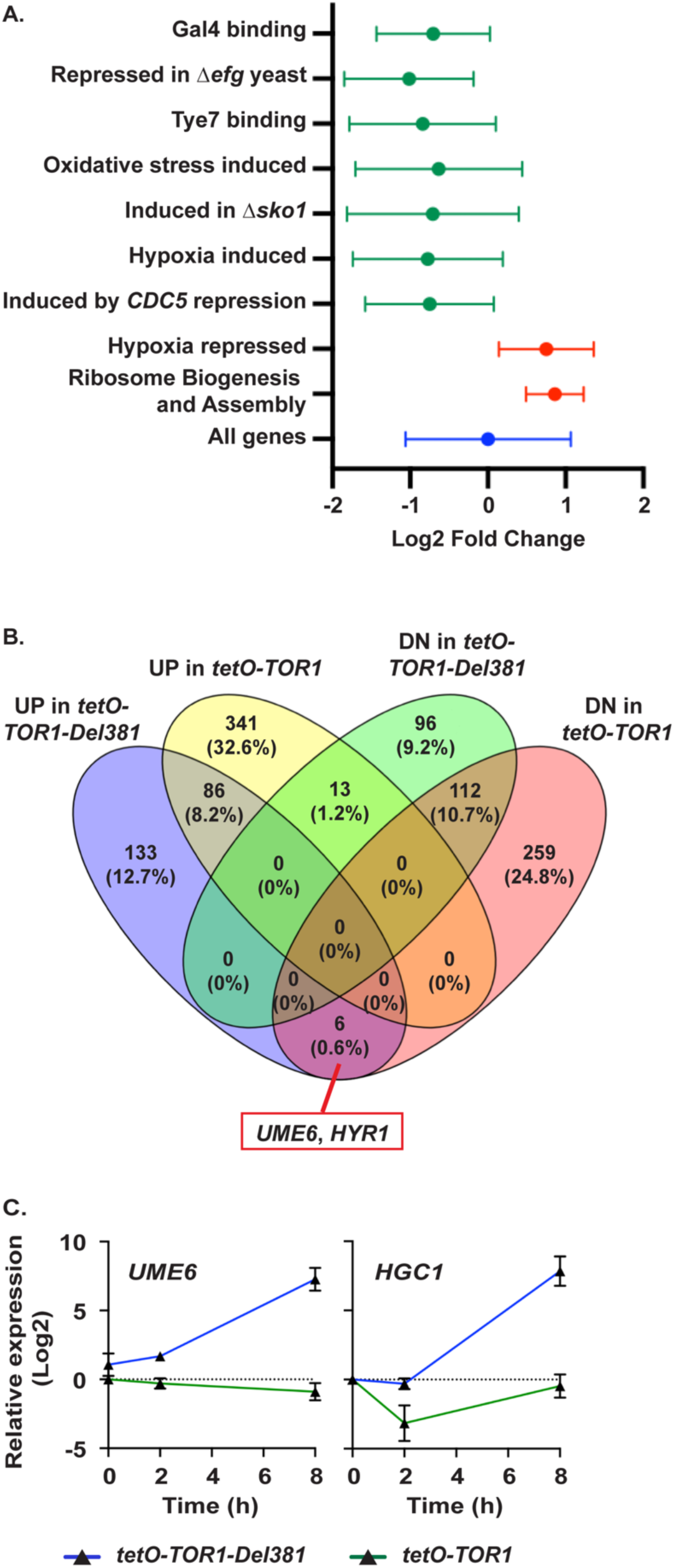
Gene expression is misregulated in cells lacking N-terminal HEAT repeats. **A.** Gene set enrichment analysis was used to identify major categories of genes significantly regulated (adjusted P value < 0.001) in *tor1/tetO-TOR1-Del381* cells (JKC1441) compared to the *tor1/TOR1* control (JKC1347). Graph shows the fold change (Log2) of genes in each indicated category in Del381 cells relative to *tor1/TOR1* cells. Full results in supplementary data. **B.** Comparison of gene expression changes (≥2-fold up and down) in Del381 (JKC1441) and FL (JKC1549) cells exposed to doxycycline for 8 h. Six genes induced in Del381 cells were suppressed in FL cells, including *UME6* and *HYR1*. **C.** Relative expression of *UME6* and *HGC1* in Del381 (JKC1441) and FL (JKC1549) cells in YPD and during exposure to 30 µg/ml doxycycline for 2 h and 8 h. Gene expression levels were compared with those in *TOR1* heterozygous cells (*tor1/TOR1*, JKC1347).

Cell wall biosynthetic gene expression of Del381 cells was perturbed, with reduced expression of genes involved in the production of cell wall sugar polymers chitin and glucan (GO:71555; Supplementary File 2), possibly contributing to these cells’ hypersensitivity to cell wall stress. In line with increased expression of hypoxia-induced genes [87], increased expression of genes involved in ergosterol biosynthesis (*ERG1, ERG3, ERG25*) was observed during repression of *TOR1-Del381* (Supplementary File 3), possibly related to increased endurance of fluconazole exposure. Additionally, Del381 cells had a 3-fold increase in expression of the fluconazole efflux pump encoding gene *MDR1* (Supplementary File 3).

Next we compared the effects of doxycycline suppression on Del381 and on FL cells (Supplementary Files 3-6). In general, repressing the *TOR1-Del381* allele with doxycycline (30 µg/ml) increased the number of differentially regulated genes identified previously, while GSEA analysis indicated that the processes affected following doxycycline treatment were similar to those perturbed in these cells without doxycycline (Supplementary Files 1-4). Many similar processes were affected in FL cells following doxycycline suppression, including ribosome biogenesis, hypoxia and oxidative stress (Fig. 10B).

Expression of the genes encoding inducers of hyphal growth, the transcriptional regulator Ume6 [89] and the hypha-promoting cyclin-like protein Hgc1 [90], was abnormally increased in Del381 cells during strong *tetO* repression (Fig. 10C). This finding may help explain the hyperfilamentous phenotype exhibited by Del381 cells under specific conditions, i.e. during anaerobic growth on acidic medium (Fig. 9A). Expression of both *UME6* and *HGC1* decreased following suppression of the *TOR1-FL* allele (Fig. 10C). Uniquely, Del381 cells exhibited increased expression of the hyphal wall protein-encoding gene *HYR1* in addition to the filamentation-inducing transcription factor *UME6* (Fig. 10B), as well as cell wall protein-encoding genes *ALS2, ALS4, ALS6* and *HWP1,* which may explain these cells’ hyperaggregation phenotype (Fig. 9B).

We further examined the capacity of the Del381 strain to mount a protective transcriptional response to oxidative stress induced by plumbagin. Exposure of wild type *C. albicans* cells to plumbagin induced a transcriptional program indicative of a strong oxidative stress response following 60 min plumbagin exposure relative to cells treated with DMSO alone (Supplementary file 7). When the responses of the Del381 and wild type cells were compared, gene categories associated with stress and environmental responses were more strongly induced in the wild type, indicating that N-terminal HEAT repeats of Tor1 were required for effective transcriptional responses to oxidative stress (Fig. 11; Supplementary Files 7, 8). These results were consistent with Del 381 cells’ hypersensitivity to oxidative stress (Fig. 4B). In the response to this important host-derived stress, Tor1 N-terminal HEAT repeats were therefore required both at the level of stress signaling and at the transcriptional level

**Fig. 11.**
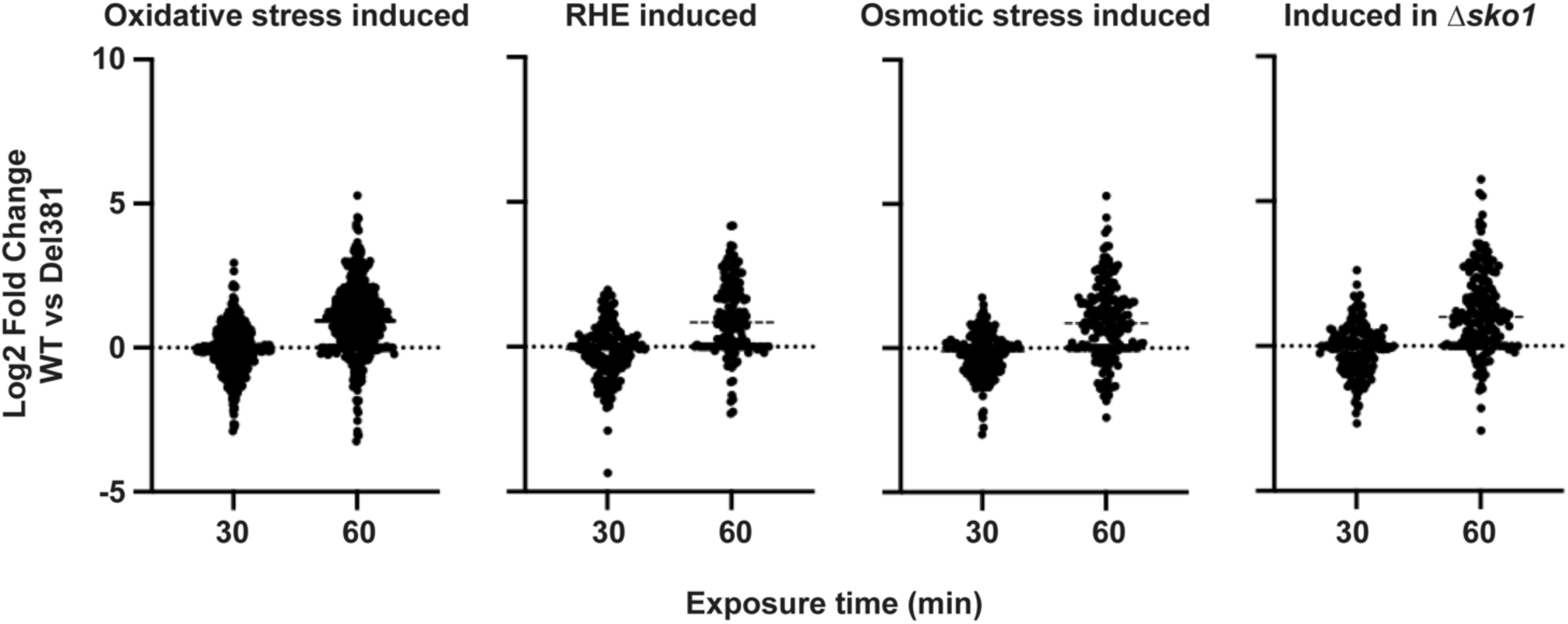
Specific gene categories are altered in cells lacking N-terminal HEAT repeats during superoxide stress. Comparison of gene expression in wild type (WT, SC5314) relative to Del381 (JKC1441) in the presence of plumbagin (10 µM) after 30 and 60 minutes. Gene categories shown were significantly represented (Q < 0.001) among genes exhibiting increased expression in the wild type relative to Del381 cells following 60 minutes incubation in plumbagin. Gene categories are those identified by Enjalbert et al. (oxidative and osmotic stress;[66]), Rauceo et al. (Δ*sko1*; [112]) and Spiering et al. (Reconstituted human epithelium induced; [113]).

## Discussion

We set out to examine whether specific roles could be assigned to a region of Tor kinase that is highly divergent between fungal and human cells, comprising the 8 most N-terminal HEAT domains. Cells whose only *TOR1* allele was transcribed from repressible *tetO* revealed specific functions to which this region contributes, when phenotypes of cells expressing an N-terminally truncated predicted protein were compared to those expressing predicted full length Tor1. While we did not measure protein levels of Tor1 as we lacked a Tor1-specific antibody and do not yet have a functional epitope tagged Tor1, qRT-PCR experiments showed that mRNA levels of both *TOR1* alleles correlated with the concentrations of doxycycline used to repress *tetO* (Fig. S1A). Similarly, growth of both *tetO*-controlled *TOR1* genotypes correlated with doxycycline concentrations in the medium, with 1 µg/ml doxycycline providing nearly complete repression of Del381 cells’ growth, while a moderate doxycycline concentration of 300 ng/ml had a small growth-repressive effect in this medium (Fig. 2A). For many experiments going forward, we therefore chose 300 ng/ml doxycycline in order to compare phenotypes in cells that had diminished but not ablated *TOR1* expression. Overexpression of either *TOR1* allele from unrepressed *tetO* in the absence of doxycycline did not lead to TORC1 hyperactivation in rich medium (Fig. 2B), as determined from phosphorylation of ribosomal protein S6, an established *C. albicans* TORC1 activity readout [25]. A simple explanation for this finding is that Tor1 kinase depends on the other components of TORC 1 for its activity, since presumably many of its activators, inhibitors and substrates are brought to their sites of physical interaction by the other complex components; overexpressing only the catalytic moiety of the complex appears to be insufficient to hyperactivate the complex (Fig. 2B).

A major input activating TORC1 signaling is availability and quality of nitrogen sources. Lack of N-terminal HEAT repeats had little impact on cells’ growth rates in the non-preferred nitrogen source proline, in the absence or presence of a moderate repressing doxycycline concentration (Fig. 2C). However, in two preferred nitrogen sources, glutamine and ammonium sulfate, cells with Tor1 lacking N-terminal HEAT repeats grew more slowly, and during moderate *tetO* repression grew only to low densities (Fig. 2C). Del381 cells similarly failed to maximize growth rates during optimal provision of phosphate- and carbon sources (Figs. 2E and 3A). These findings together indicated that the TORC1 response to favorable nutritional conditions, to coordinate anabolic processes like translation and DNA replication with provision of metabolic intermediates and harvesting of energy from carbon sources, depends on at least one function provided by the N-terminal HEAT repeats comprising amino acid residues 1-381. Since this failure occurred in response to three central macronutrients, nitrogen-, carbon- and phosphate sources, we speculate that the defective activity of N-terminally truncated Tor1 relates not to defective sensing of individual nutrients, but to misregulation of a pro-anabolic output that normally results from TORC1’s integration of favorable nutritional inputs together with absent perception of unfavorable stressors. In the host, inability to take advantage of favorable nutrient conditions will place a member of the mucous membrane microbiome at a distinct disadvantage to other organisms that maximize their growth rates when nutritional conditions permit. The ability to optimally accelerate growth during favorable nutrient conditions is therefore a critical role of TORC1 for *C. albicans*.

While Del381 cells were unable to appropriately accelerate growth in optimal conditions, they inappropriately activated TORC1 signaling when nutrient conditions were suboptimal, as indicated by an excessive P-S6 signal in the nonpreferred nitrogen source proline and in a lower glucose concentration of 0.25% (Figs. 2D and 3D). Rps6 phosphorylation was not simply indiscriminately activated, since Del381 cells growing in the absence of glucose (without an added carbon source) or in the nonfermentable carbon source glycerol, had no detectable P-S6 signal, in line with wild type and FL cells (Fig. 3D). Hence Del381 cells seemed to be lacking a fine-tuning function of TORC1 activity, while “on” and “off” switches remained functional.

Rps6 phosphorylation, whose biochemical function remains unknown [91], corresponds to translational activation in *C. albicans* [25]. If Del 381 cells inappropriately activated translation in suboptimal nutrient conditions, their slower growth may have been due to futile protein production with wasteful consumption and premature exhaustion of scarce resources. We observed slower oxygen consumption of cells overexpressing *TOR1-Del381* compared to those overexpressing *TOR1-FL*, or to wild type cells, during growth in glucose (Fig. 3E), suggesting that energy extraction of these cells from glucose was not coordinated with translation. To examine translational control in these cells from another aspect, we assayed the behavior of Gcn2 kinase through the phosphorylation state of its target, eIF2*α*. Strikingly, Gcn2 appeared to be hyperactive in Del381 cells in most conditions (except for cell wall stress) compared to wild type and FL cells: eIF2*α* was hyperphosphorylated, indicating inhibition of translation initiation by Gcn2, in conditions where Del381 cells’ P-S6 signal was hyperintense indicating excessive translation activation e.g. during growth in proline (Fig. 2D). (Hyperphosphorylation of eIF2*α* additionally occurred in Del381 cells in conditions where the P-S6 signal was undetectable e.g. in stationary phase and in medium without an external carbon source or with glycerol (Figs. 2D and 3D)). Experiments we then performed with these cells during oxidative stress may have shed some light on this paradox, as described below.

We examined intracellular ROS of Del381 cells compared with FL and wild type cells, since slow oxygen utilization of these cells (Fig. 3E) suggested that their putative decreased mitochondrial activity might generate less intrinsic ROS, as was indeed the case (Fig. 4A). This alteration did not correspond to increased tolerance of extrinsic oxidative stress, however: to the contrary, Del381 cells were strongly hypersensitive both to peroxide- and to superoxide anion-stress, as induced by H_2_O_2_ and plumbagin, respectively (Fig. 4B). Their oxidative stress signaling through the Hog1 kinase was weak (Fig. 4C), and their transcriptional oxidative stress responses were blunted (Figs. 10A and 11). During invasion of the host, *C. albicans* is exposed to ROS from host phagocytes [49, 52, 92], hence an activity of Tor1 and its N-terminal HEAT repeats in coordinating oxidative stress responses with HOG pathway signaling is predicted to be indispensable for the host interaction.

A rapid response to oxidative stress in the model yeasts *S. cerevisiae* and *S. pombe* as well as in *C. albicans* is widespread inhibition of translation initiation and elongation, while stress responses are mobilized [68, 93–95]; we therefore examined Del381 cells’ Rps6 phosphorylation during oxidative stress (plumbagin exposure). The P-S6 signal of Del381 cells was hyperintense in the first 20 minutes of plumbagin exposure (Fig. 4D), indicating inappropriate translation activation. At the same time, translation inhibition through Gcn2 activity was more active than in wild type or FL cells, as assayed by eIF2*α* phosphorylation (Fig. 4D). Since the final common effector of these two signaling pathways is the activity of the translational machinery, we examined which of the two pathways that gave opposing signals - TORC1 an activating- and Gcn2 an inhibitory signal - had the decisive effect, or the “last word.” We examined translation of a heterologous message encoding GFP, which had no known intrinsic regulatory motifs responding to oxidative stress, transcribed from regulatable promoters *tetO* or *pMAL2*. In wild type cells, translation of a GFP message transcribed from inducible *tetO* was delayed during oxidative stress exposure (Fig. 5A). GFP expressed from *pMAL2* during oxidative stress was translated inappropriately early in Del381 cells compared with wild type, *tor1/TOR1* heterozygous, and FL cells (Fig. 5B), indicating that hyperactivation of translation by TORC1 prevailed over translation inhibition by Gcn2.

Why was translation inhibition through Gcn2 upregulated in the same cells (we examined the same protein extract samples for these comparisons of P-S6 and P-eIF2*α* signal intensities) in which translation was hyperactivated as assayed through P-S6? Since in *S. cerevisiae*, Gcn2 directly phosphorylates the N-terminus of Kog1 [96] to downmodulate TORC1 activity, we surmised that the missing HEAT repeats in Del381 cells normally participate in downregulation of translation by Gcn2 through Kog1 during oxidative stress; when their activity is absent in these cells, Gcn2 develops compensatory hyperactivity. Excessive GFP translation during oxidative stress in these cells showed that this compensatory upregulation of eIF2*α* phosphorylation is not sufficient, in the end, to suppress hyperactive translation in Del381 cells.

However, hyperactive translation did not invariably predict growth defects: Del381 cells exposed to the plasma membrane stressor SDS had inappropriately high levels of P-S6 while enduring this stress well when their *TOR1-Del381* allele was not repressed (Fig. 8A-C). Strikingly, during *TOR1-Del381* overexpression, these cells showed no growth defect in fluconazole, and even during moderate *tetO* repression their growth defect in this drug was minor compared to their defect in vehicle, indicating enhanced fluconazole endurance (Fig. 8D). These cells also endured amphotericin B exposure well while their *TOR1-Del318* allele was overexpressed (Fig. 8D). Conversely, growth defects of Del381 cells in a specific stress condition were not strictly tied to defective signaling of the cognate stress response pathway, as exemplified by phosphorylation of the cell wall integrity pathway component Mkc1 (Figs. 6C, S4A). Instead these growth defects may be caused by inefficient respiration (Fig. 3E) and hence an inadequate ATP supply for biosynthesis of cell wall polysaccharide monomers, i.e. nucleotide sugars [97]; additionally, transcriptional induction of cell wall polysaccharide biosynthesis was deficient in Del381 cells (Supplementary File 2). It is also possible that stress response circuits between TORC1 and Rho1 are disrupted by a conformational change of TORC1 at the site of the Kog1 domain bound alternatively by Tap42 phosphatase or by Rho1 [75, 98, 99]. Rho1 directs responses to heat and other stressors as well as to cell wall stress in this circuit [75], and its perturbation may not necessarily be detectable as defective Mkc1 activation [75]. Taken together, our findings preclude simplistic interpretations of the role of Tor1 N-terminal HEAT repeats based only in their repression of inappropriate translation. They highlight that TORC1 inputs and outputs from this region of Tor1 inevitably interact during regulation of manifold processes that respond to combinations of environmental conditions to which the fungus is exposed, as shown in Table 1. In the experiments we report here, single stressors were examined in isolation; actually during infection, combinations of these stressors act on *C. albicans* as often emphasized by Brown and colleagues e.g. in [100].

**Table 1.** Del381 cell responses to specific stress types.

| Table 1. Del381 cell responses to specific stress types. |  |  |  |
| --- | --- | --- | --- |
| Stress type | Hyperintense P-S6 | Growth defect | Defective specific pathway signaling |
| Non-preferred nitrogen source | yes | no | not tested |
| Low glucose | yes | no | not tested |
| Low inorganic phosphate | yes | no | not tested |
| Oxidative stress | yes | yes | yes (Hog1 phosphorylation) |
| Cell wall stress | no | yes | no (Mkc1 phosphorylation) |
| Heat stress | no | yes | not tested |
| Plasma membrane stress | yes | no | not tested |

Of particular interest will be investigation of roles of N-terminal HEAT repeats in metabolic regulation. TORC1 inhibition by rapamycin is known to change the metabolome within minutes, suggesting that TORC1 regulates metabolic enzymes by posttranslational modification [101]. When some signals to TORC1 cannot be received and others from TORC1 cannot be sent because the interaction domains for relevant up- or downstream partners are absent, specific metabolic derangements would be expected. In addition to decreased oxygen consumption of Del381 cells (Fig. 3E), our transcriptional analysis also shows perturbation of carbon source metabolism in Del381 cells.

Del 381 cells’ inability to grow on lactate is notable. They grew well on other non-fermentable carbon sources (Fig 3B), indicating that defective growth on lactate is not simply attributable to defective respiration. These cells also grew well on medium of pH2 with a fermentable– or non-fermentable carbon source (Fig. S2), suggesting that a more specific defect than acid stress intolerance is responsible for this phenotype. Transcription of the cell-surface lactate transporter-encoding gene *JEN1* [102] was not decreased in Del381 cells (Supplementary File 2). Expression of the lactate dehydrogenase gene *CYB2*, known to be required for *Candida glabrata* virulence in an insect model [103], was also not down- but actually moderately upregulated (to a 1.37 Log2 fold change) in Del381 cells compared with *tor1/TOR1* cells (Supplementary File 1). Use of lactate as the carbon source is known to affect *C. albicans’* cell wall composition and interaction with the host [59–61]. In *S. cerevisiae*, expression of the human lactate dehydrogenase isoform LDHB, which preferentially converts lactate to pyruvate, increases lactate tolerance [104]. A wide array of processes contributes to lactate tolerance in this model yeast as recently reviewed by Thevelein and colleagues [105]; while the extent of conservation of these processes between *S. cerevisiae* and *C. albicans* is not clear, divergence of mitochondrial biogenesis and mitochondrial protein import between these species with implications for lactate metabolism has been demonstrated [106]. How these processes relate to an involvement of Tor1 N-terminal HEAT repeats in tolerance of lactate and in respiration is currently not known; a better understanding may emerge from analysis of Del381 cells’ metabolism.

We focused on TORC1 though truncation of Tor1 is likely to also affect its role in the rapamycin-insensitive TOR complex 2 (TORC2) [107]. The prominence of inappropriately active anabolic processes in Del381 cells during oxidative stress, like ribosome biosynthesis and translation, indicates that Tor1 N-terminal HEAT repeats in stress responses we examined, act in the context of TORC1. Which TORC2 functions are disrupted by loss of Tor1 N-terminal HEAT repeats will need to be studied in further investigations of this complex’s interaction partners.

Our findings highlight the importance of intact, physiologically modulated TORC1 signaling in specific stress responses critical to *C. albicans*’ interaction with the human host (Fig. 12). Suboptimal TORC1 reactivity hence is predicted to impair *C. albicans*’ virulence; differences in their TORC1 signaling system were previously discussed as underlying the virulence differences between the frequent pathogen *C. albicans* and its close relative, the rarely invasive *C. dubliniensis* [108]. Our findings also demonstrate that stepwise functional dissection of distinct segments among the large number of interaction domains of the Tor1 kinase is possible. Together with identification of interaction partners for each segment, as well as changes in the phosphoproteome and metabolic shifts during perturbation of distinct segments, the anatomy of the afferent and efferent pathways to each element of the “brain of the cell” can be defined in more detail than has so far been undertaken. Small molecules that disrupt some of these pathways could become “nonimmunosuppressive rapamycin analogs” as envisioned by Cruz et al. [6]. In addition to an approach that targets the highly conserved Tor kinase domain in a more fungal-selective manner [6], the substantial structural differences between human and fungal TORC1 components including the Tor kinase itself (Fig. 1) could be exploited for fungal-specific targeting of essential functions, particularly those required during the host interaction.

**Fig. 12.**
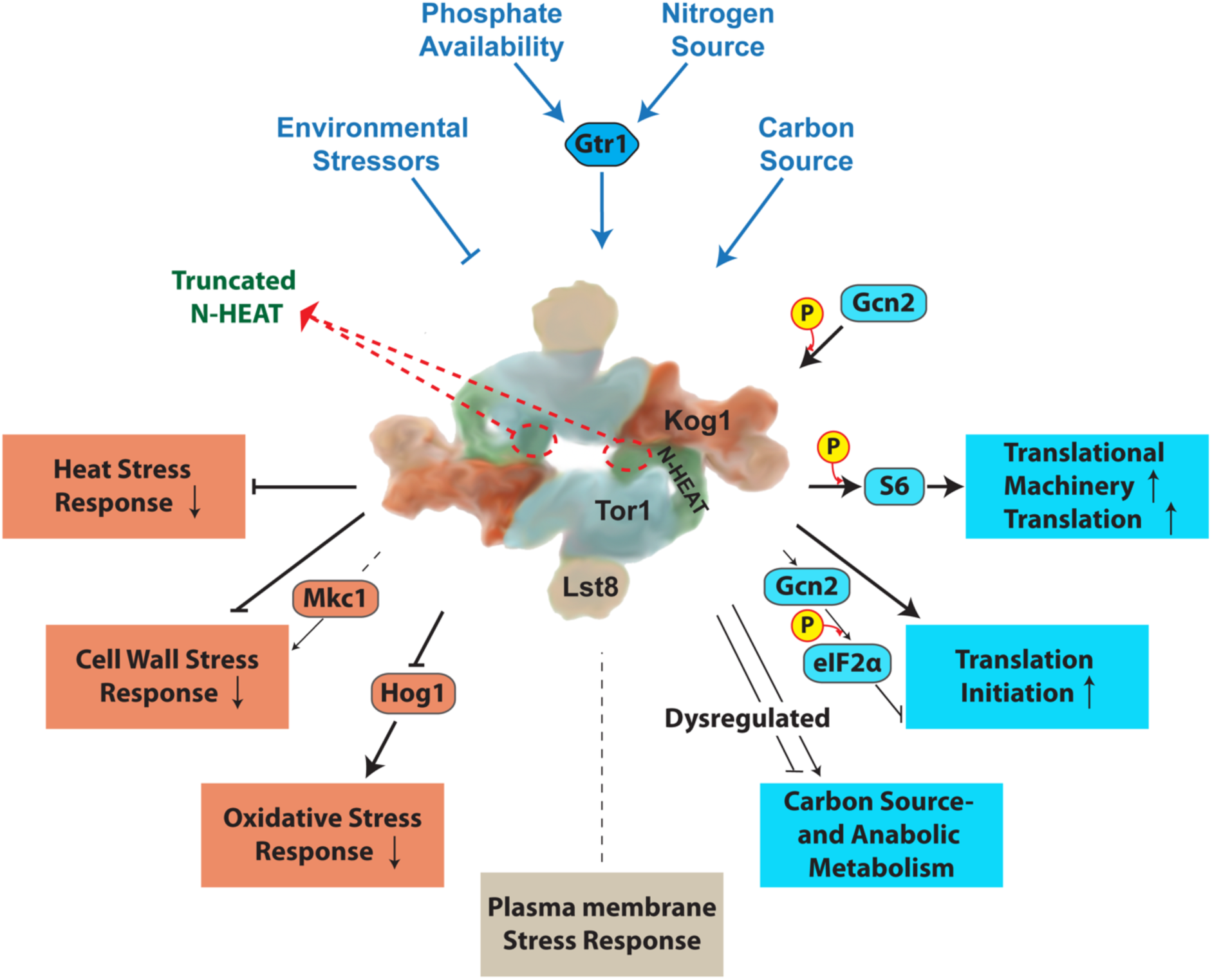
Model of the role of Tor1 N-terminal HEAT repeats in nutritional- and stress responses. Cartoon of TORC1 without Tco89 after PDB:6BCX and [14] with regulatory inputs and outputs examined in this work. Tor1 N-terminal HEAT repeats are shown in green and the remainder of the protein in blue.

## Methods

All experiments were performed in at least 3 biological replicates on different days unless otherwise stated. Graphs, blots and images show a representative experiment of the replicates, except where indicated that data from 3 biological replicates were combined into a graph.

### Strains and culture conditions

*C. albicans* strains used are shown in Table S1. Strains were constructed as described in [42], with plasmids shown in Table S2 and oligonucleotides shown in Table S3, using sequences obtained from the Candida Genome Database [109]. To minimize phenotypic artifacts originating from genomic events unrelated to the targeted introduced mutations, all genotypes examined were constructed from at least 2 independently engineered heterozygous strains. *C. albicans* cells were grown as described in [42] unless otherwise stated.

### Growth curves

Stocks stored at -80° were recovered on YPD agar medium for 2 days. Cells were scraped from the plate and washed once in 0.9% NaCl and diluted at OD_600_ 0.01 (for phenotypes for antifungal sensitivity) or OD_600_0.1 (for nutrient phenotypes) in 150 µl medium in flat bottom 96-well dishes. OD_600_ readings were obtained every 15 minutes in a BioTek^TM^ Synergy^TM^ 2 Multi-Mode Microplate Reader (Winooski, VT, USA). Standard deviations of 3 technical replicates, representing separate wells, were calculated and graphed in Graphpad Prism Version 9.1.0 (216).

### Western Blot

Cell lysis and Western blotting were performed as described in [25]. Antibodies used are listed in Table S4. For densitometry, ImageJ (imagej.net/welcome) software (opensource) was used to quantitate signals obtained either from KODAK Image Station 4000MM or from Azure biosystems c600.

### Growth of cell dilution spots on solid media

Serial cell dilution spotting assays on agar media was performed as previously described [42]. Briefly, cells recovered from glycerol stock were grown on YPD agar media for 36-48 h, then washed in 0.9% NaCl and diluted with 0.9% NaCl in 5-fold steps from a starting OD_600_ of 0.5 in a microtiter plate, then pin transferred to different agar media. Plates were imaged after 48 hours of incubation at 30° unless stated otherwise.

### Oxygen consumption measurement

Oxygen consumption was measured using a Clark-type electrode (dual digital-model 20; Rank Brothers. Ltd., Cambridge, United Kingdom) at 25°, according to the manufacturer’s instructions. *C. albicans* cells were grown to mid-logarithmic phase in YPD at 30°, washed twice in 0.9% NaCl and resuspended in the same solution. Cell suspensions were prepared at 5×10^6^ cells/ml. As a control for oxygen saturation, one chamber was filled with 700 μl 0.9% NaCl; in the second chamber 650 μl of cell suspensions was added to 50 μl glucose (to achieve a 2% final glucose concentration). Oxygen saturation was recorded every 3 minutes.

### ROS measurement

Yeast cells grown overnight in YPD for 15 hours were washed twice with 0.9% NaCl and diluted in SC medium (LoFlo) at OD_600_ of 0.5. The fluorescent dye 2’7’-dichlorodihydrofluorescein diacetate (DCFDA) (Sigma, Cat#D6883) was added into the medium to a final concentration of 50 μM. After incubation for 90 minutes, cells were washed twice with 0.9% NaCl and 100 μl per well was placed in a 96-well black plate with optical bottom (Thermo Scientific Nunc), in triplicate. The intensity of fluorescence was read in a BioTek^TM^ Synergy^TM^ 2 Multi-Mode Microplate Reader (Winooski, VT, USA) at excitation 485 nm and emission wavelength 528 nm, and a ratio of fluorescence intensity to OD_600_ of the culture was calculated.

### Hyphal morphogenesis assay

Cells were revived from frozen stocks on solid YPD medium for 2 days, washed and resuspended in 0.9% NaCl to OD_600_ of 0.1. Variations between single colonies and colony density effects were minimized by spotting 3 μl cell suspension at 6 equidistant points, using a template, around the perimeter of an agar medium plate. M199, YPD plus 10% serum and Spider medium were used. M199 medium was buffered to pH 5 with 100 mM MES and to pH 7 with 165 mM MOPS. Plates were incubated at 37° between 2 and 4 days. For phenotypes under anaerobic conditions, plates were incubated in an anaerobic chamber (COY lab products) with a mix of 10% Hydrogen, 10% Carbon dioxide and Nitrogen balance at 37°. All panels shown represent at least 2 biological replicates.

### Flocculation

Cells from YPD plates with 10 ng/ml doxycycline were collected and washed once in 0.9% NaCl, then inoculated into YPD medium containing 100 ng/ml doxycycline with a starting OD_600_ of 0.2. Cells were grown at 30° till exponential phase (∼4 h). Cells were washed twice in 0.9% NaCl and inoculated into 5.5 ml Spider medium (with additional 0.3 mM histidine) with or without 300 ng/ml doxycycline, with final OD_600_ of 0.7. Cultures were incubated at 37°, 200 rpm for 3 h, then settled at room temperature for 15 min before imaging.

### Quantitative real-time PCR analysis

For *tetO*-repression and -derepression, cells were grown in YPD with 2 µg/ml doxycycline with starting OD_600_ of 0.7 for 4 h (Time 0), washed twice with 0.9% NaCl and inoculated into YNB without ammonium sulfate ((NH_4_)_2_SO_4_) supplemented with 10 mM Proline (Pro) as sole nitrogen source, at starting OD_600_ of 0.5, cells were collected at 45 min and 90 min time points. For plumbagin treatment, cells were grown in YPD with 5 ng/ml doxycycline with starting OD_600_ of 0.3 for 3.5 h (Time 0), washed twice in YPD with 5 ng/ml doxycycline and inoculated into YPD with 5 ng/ml doxycycline and 10 µM plumbagin, at starting OD_600_ of 0.4. Cells were collected at 20 min and 40 min time points centrifugation at the indicated time-points, washed 2 times in ice-cold normal saline (0.9% NaCl) and re-suspended in 1 ml cold TRI Reagent (Molecular Research Center Inc, TR118). Total RNA was extracted using Direct-zol RNA Miniprep Plus kit (Zymo Research, R2073) and reverse transcription was performed using qScript cDNA Synthesis Kit (Quanta Bio, 95047-100), both according to the manufacturer instructions. Quantitative real-time PCR was performed with Maxima SYBR Green/ROX qPCR Master Mix (2X) (ThermoFisher, K0222) with *CaTOR1* gene specific primers, using QuantStudio-3 96-well 0.2 ml Block (ThermoFisher). Gene expression levels were measured relative to *C. albicans ACT1* expression and q-PCR data were analyzed with QuantStudio.

### Transcript profiling

Transcriptomes of JKC1549 (*tor1/tetO-TOR1*) and JKC1441 (*tor1/tetO-TOR1-Del381*) were compared with those of the heterozygous control JKC1347 (*tor1/TOR1*). Each strain was cultured for 24 h on solid YPD containing 5 ng/ml of doxycycline. Cells were harvested from plates, washed twice in phosphate buffered saline and inoculated into 50 ml of YPD to an OD_600_ of 0.1 in the absence or presence of doxycycline (30 µg/ml). Cells were harvested after 2 h and 8 h growth. Cell pellets were flash frozen in liquid N_2_ and RNA extraction was performed using the Qiagen RNeasy mini-kit as described [110]. The microarrays used in this study were designed from assembly 21 of the *C. albicans* genome using eArray from Agilent Technologies (design ID 017942) [111]. A total of 6,101 genes (including 12 mitochondrial genes) are represented by two sets of probes, both spotted in duplicate. Total RNA (100 ng) was labelled with Cy5 or Cy3 using the Two-Color Low Input Quick Amp labelling Kit (Agilent Technologies) according to the manufacturer’s instructions. Array hybridization, washing, scanning and data extraction was carried out as described by Haran et al. [110]. Each condition was examined with four biological replicates including dye swapped replicates. Data for each feature was background corrected and expression normalized using the Lowess transformation method in GeneSpring GX13 (Agilent Technologies). Differential expression between pairs of conditions was examined by comparing relative expression ratios (Log2 values) among genes that satisfied a post-hoc test (Storey with Bootstrapping) with a corrected Q value *≤*0.05. Gene set enrichment analysis (GSEA) was carried out as described by Haran et al. [110]. The full data set can be accessed at the NCBI GEO repository (Accession GSE182186).

To compare the transcriptional responses of JKC1361 (wild-type), JKC1549 (*tor1/tetO-TOR1*) and JKC1441 (*tor1/tetO-TOR1-Del381*) to plumbagin, cells were grown as above and used to inoculate 200 ml YPD supplemented with 5 ng/ml doxycycline at OD_600_ of 0.3. These cells were grown for 3.5 hours, harvested and used to inoculate a fresh 50 ml YPD at OD_600_ of 0.4. These cultures were treated with plumbagin (10 µM) and incubated at 30° at 200 rpm and RNA was harvested at 30 min and 60 min incubation, as described above. A wild-type JKC1361 culture was also treated with DMSO alone (50 µl) in a control experiment. Three biological replicates were generated for each time point and RNA samples were pooled for cDNA sequencing Using Oxford Nanopore Technologies (Oxford Nanopore Technologies, Oxfordshire, UK) PCR-cDNA Sequencing Kit (SQK-PCS108) and barcoding kit (SQK-LWB001) according to the manufacturer’s instructions. Briefly, total RNA (50 ng) was reverse transcribed to cDNA using the PCR-cDNA Sequencing Kit (Oxford Nanopore Technologies) and SuperScript IV reverse transcriptase (Carlsbad, California, United States) according to the Oxford Nanopore Technologies protocol (version PCB_9037_v108_revF_30Jun2017). Following this, cDNA was amplified and indexed using the PCR Barcoding Kit (Oxford Nanopore Technologies) and LongAmp Taq 2X master mix (New England Biolabs). The resulting cDNA libraries were sequenced on a MinION (MIN101B) device with a FLO-min106 SpotON R9.4 Flowcell using the MinKNOWN software v1.7.10 (Oxford Nanopore Technologies). Fast5 reads that passed filtering where basecalled and demultiplexed using the Albacore software v2.2.3 (Oxford Nanopore Technologies). Porechop v0.2.4 (https://github.com/rrwick/Porechop) was used to further trim reads and a total of 5.16 million reads were obtained. Sequence reads were aligned to the SC5314 genome (Assembly 21) using BWA-MEM and the resulting SAM files were analysed in Strand NGS (Strand Life Sciences). Differential gene expression between pairs of conditions was examined using genes that satisfied a post-hoc test (Storey with Bootstrapping; Q *≤*0.05) using GSEA [110]. Raw data can be accessed at NCBI (Accession PRJNA753651).

### Statistical analysis

For growth curves, the total area under the curve and its standard deviation were obtained and normalized based on the mean area for the control group, within a 95% confidence interval, using XY analysis in Prism 9 GraphPad (GraphPad Software, Inc., CA, USA). Statistical significance among groups was determined using an unpaired t-test with Welch’s correction, with two-tailed P values.

## Acknowledgments

We thank Tahmeena Chowdhury Godin for foundational experiments and inspiration for this work. M.A.-Z. was funded in part by the Alfonso Martin Escudero Foundation (Madrid, Spain). P.R.F. was funded by Science Foundation Ireland Grant 11/RFP.1/GEN/304. J.R.K. was funded by NIAID R21AI096054 and NIAID R21AI137716.

## Supplementary Files containing expression data

| Supp. File Number | Content |
| --- | --- |
| 1 | Genes showing 1.5 fold expression increase in D381 relative to heterozygous *TOR1/tor1* |
| 2 | Genes showing 1.5 fold expression decrease in D381 relative to heterozygous *TOR1/tor1* |
| 3 | Genes showing increased expression in D381 following 2h exposure to Doxycycline |
| 4 | Genes showing decreased expression in D381 following 2h exposure to Doxycycline |
| 5 | Genes showing increased expression in *tetO-TOR1-FL* following 2h exposure to Doxycycline |
| 6 | Genes showing decreased expression in *tetO-TOR1-FL* following 2h exposure to Doxycycline |
| 7 | Gene expression SC5314 exposed to 10 µM Plumbagin vs SC5314 exposed to DMSO |
| 8 | Gene expression in SC5314 vs *tetO-TOR1-D381* in 10 µM Plumbagin |

## Supplementary Figure Legends

**Fig. S1.**
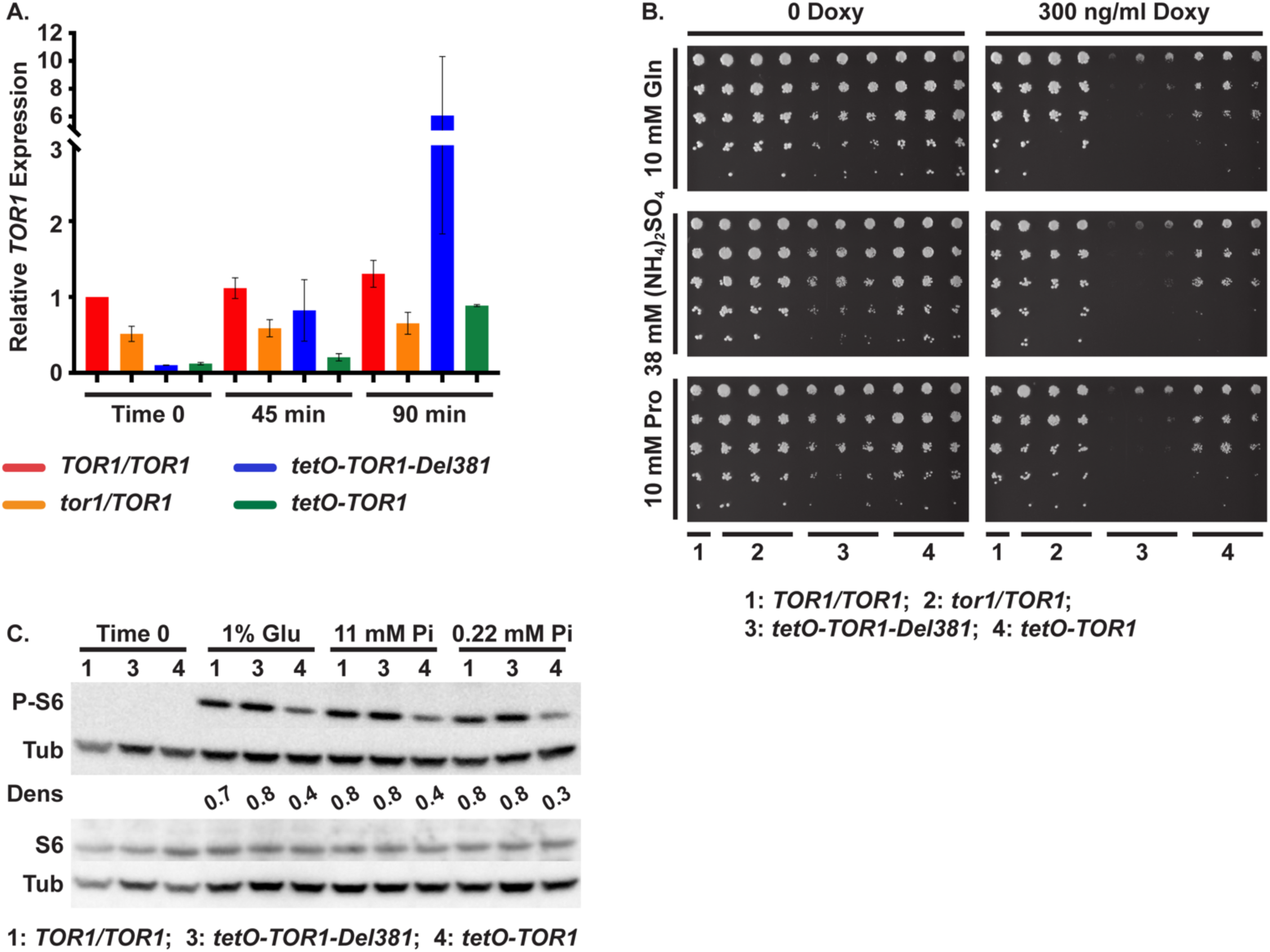
*TOR1* expression from *tetO* rebounded more rapidly in Del381 cells after release from doxycycline repression. **A. *TOR1* mRNA levels during *tetO* repression and derepression.** Cells of indicated genotypes were pre-grown in YPD medium with 2 µg/ml doxycycline for 4 h, and inoculated into YNB without ammonium sulfate supplemented with 10 mM proline (Pro) as sole nitrogen source. Cells were harvested at the end of pre-growth (Time 0) and at 45 min and 90 min after inoculation. After qRT-PCR the *TOR1* expression level was measured relative to *ACT1* expression levels. Values are expressed as means ± SD (2 biological replicates using different *TOR1* mutant lineages). (*TOR1/TOR1*, JKC1713; *tor1/TOR1*, JKC1347, JKC1346; *tetO-TOR1-Del381*, JKC1441, JKC1445; *tetO-TOR1,* JKC1549, JKC 1546). **B. Growth of cell dilutions on media containing different nitrogen sources.** Dilutions of cells of indicated genotypes were spotted onto YNB agar medium without ammonium sulfate, supplemented with 10 mM glutamine (Gln), 38 mM ammonium sulfate ((NH_4_)_2_SO_4_) or 10 mM proline (Pro) as sole nitrogen source, without or with 300 ng/ml doxycycline (300 Doxy). (*TOR1/TOR1*, JKC1713; *tor1/TOR1*, JKC1345, JKC1346, JKC1347; *tetO-TOR1-Del381*: *tor1/tetO-TOR1-Del381*, JKC1442, JKC1445, JKC1441; *tetO-TOR1*: *tor1/tetO-TOR1-FL,* JKC1543, JKC1546, JKC1549). **C. Rps6 phosphorylation during phosphate starvation or repletion.** Cells of indicated genotypes were pre-grown in YPD medium with 2 µg/ml doxycycline for 4 h, and inoculated into normal YNB supplemented with 1% glucose (1% Glu) or YNB without phosphate supplemented with either 11 mM or 0.22 mM KH_2_PO_4_ (Pi) and 2% glucose. Cells were harvested at 45 min, total protein extract was probed with antibody to phosphorylated Rps6 (P-S6), total Rps6 (S6), and tubulin (Tub) as loading control. Dens: signal intensity ratio of P-S6 to Tub. Strains *TOR1/TOR1*, JKC1361; *tor1/TOR1*, JKC1347; *tetO-TOR1-Del381*: *tor1/tetO-TOR1-Del381*, JKC1441; *tetO-TOR1*: *tor1/tetO-TOR1-FL,* JKC1549.

**Fig. S2.**
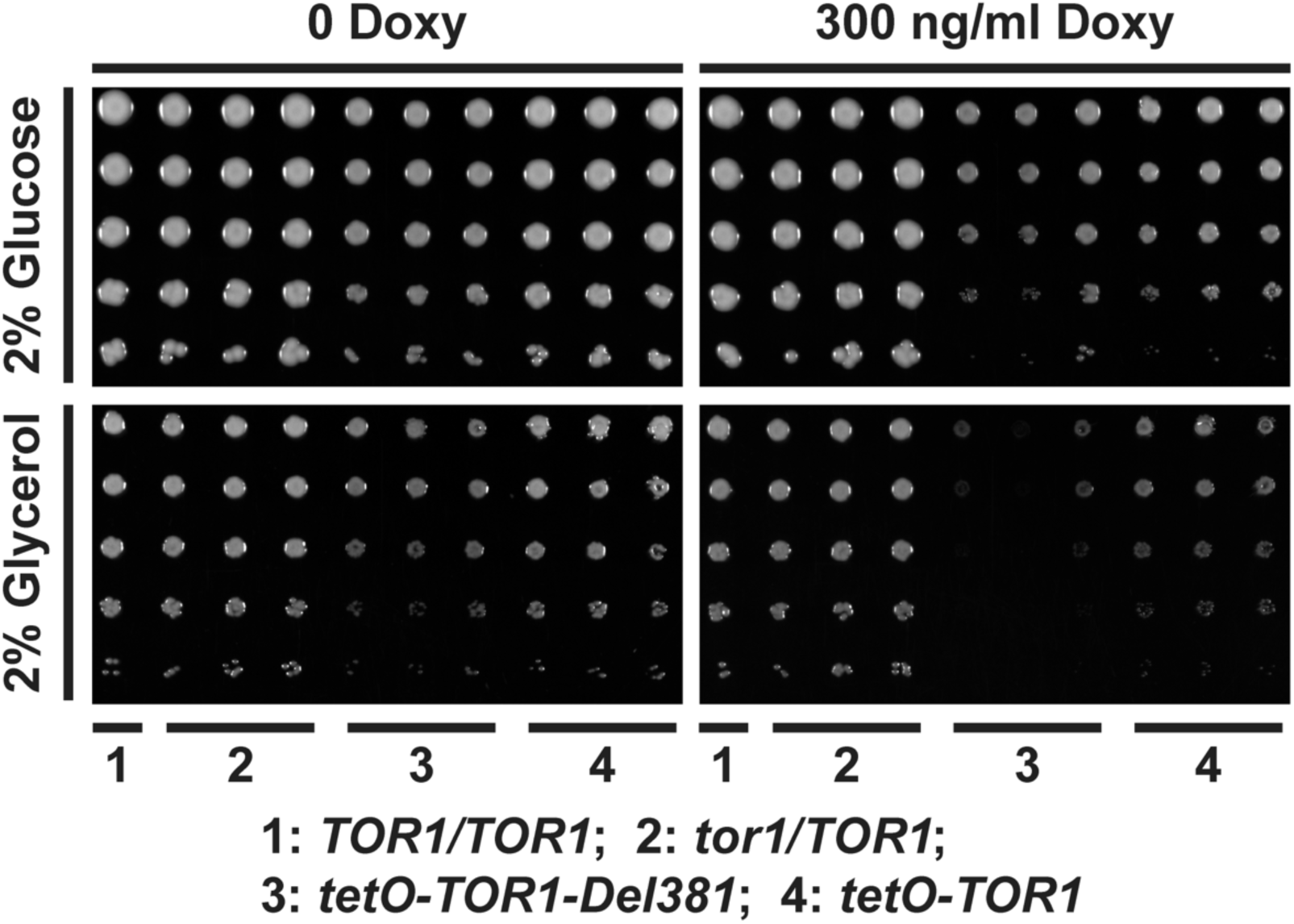
Del381 cells were able to grow at pH2 on a fermentable- and a non-fermentable carbon source. Cells of indicated genotypes were spotted onto YPD agar medium containing glucose or glyecerol (2% w/v), without or with 300 ng/ml doxycycline. Medium was buffered to pH2 with 100 mM MES. Representative of 2 biological replicates. Strains *TOR1/TOR1*, JKC1713; *tor1/TOR1*, JKC1345, JKC1346, JKC1347; *tetO-TOR1-Del381*, JKC1442, JKC1445, JKC1441; *tetO-TOR1,* JKC1543, JKC1546, JKC1549.

**Fig. S3.**
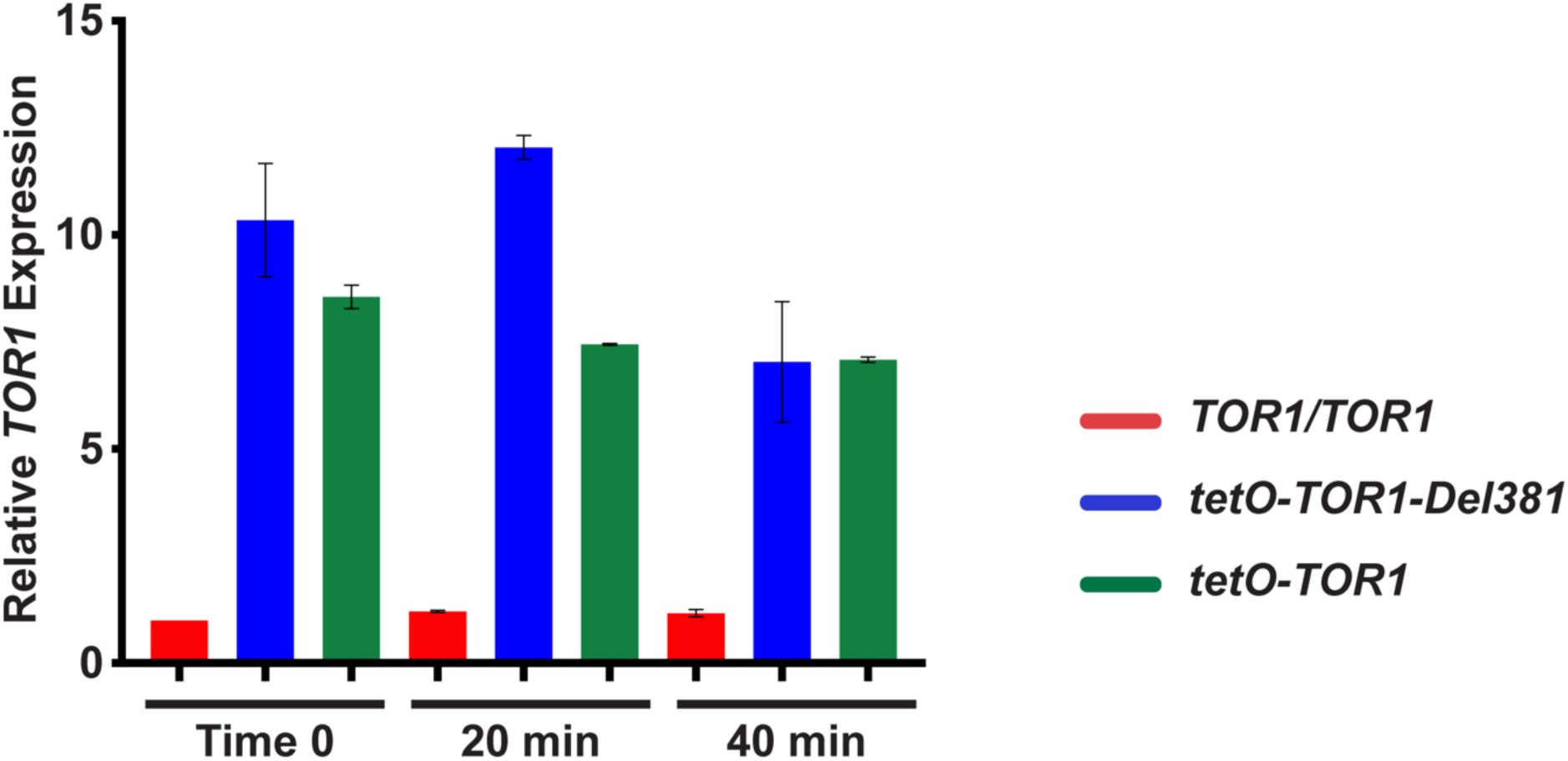
*TOR1* overexpression from *tetO* was not impaired by plumbagin treatment. Cells of indicated genotypes were pre-grown in YPD medium with 5 ng/ml doxycycline for 3.5 h, and inoculated into YPD medium with 5 ng/ml doxycycline and 10 µM plumbagin. Cells were harvested at the end of pre-growth (Time 0) and at 20 min and 40 min after plumbagin exposure. After qRT-PCR, the *TOR1* expression level was assayed relative to *ACT1* expression. Values are expressed as means ± SD (2 biological replicates using different *TOR1* mutant lineages). Strains *TOR1/TOR1*, JKC1713; *tor1/TOR1*, JKC1347, JKC1346; *tetO-TOR1-Del381*:, JKC1441, JKC1445; *tetO-TOR1,* JKC1549, JKC 1546.

**Fig. S4.**
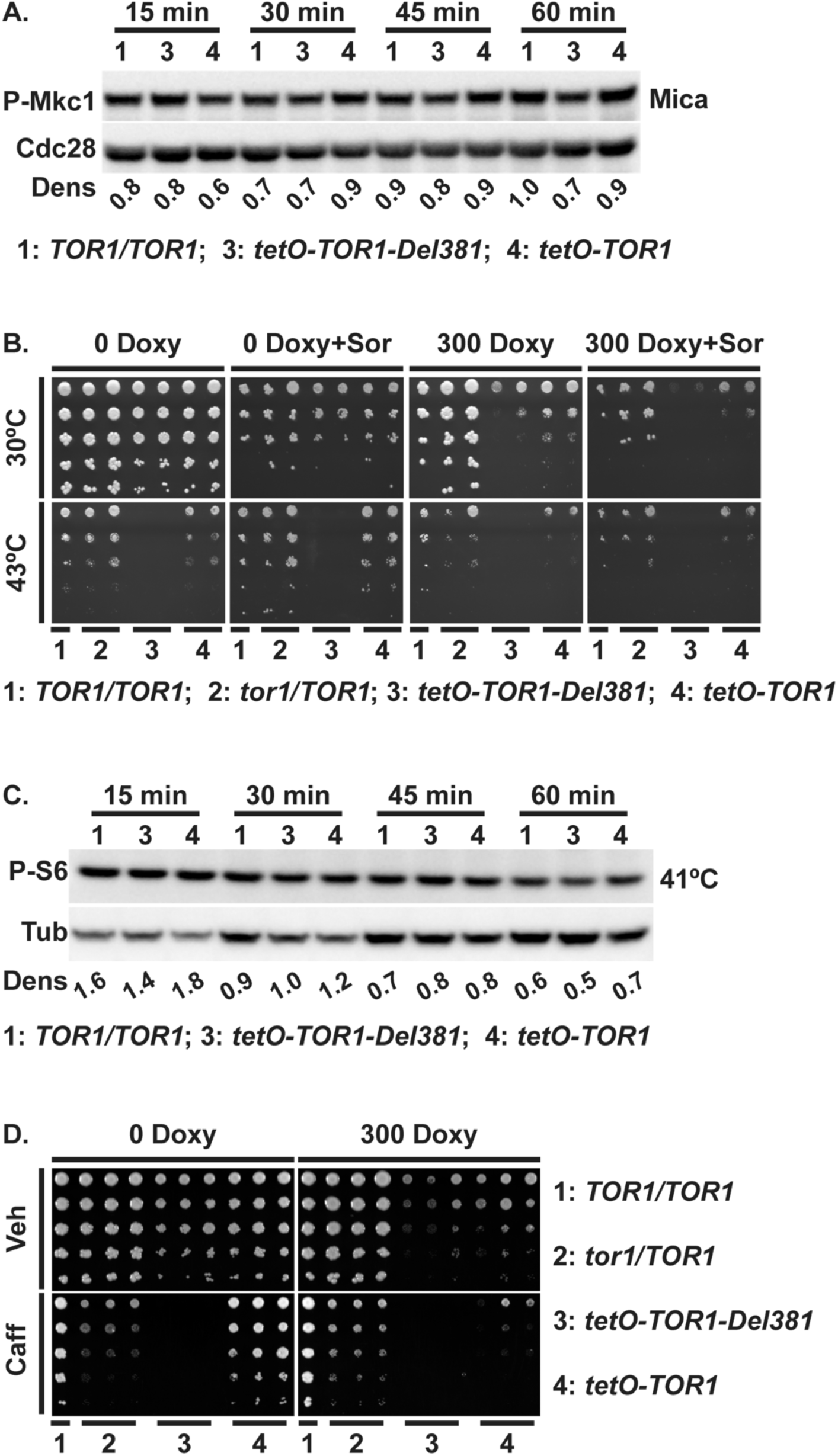
Del381 cells, hypersensitive to heat- and cell wall stress, were not defective in cell wall integrity signaling while heat stress did not affect P-S6 intensities. A. Rps6 phosphorylation during cell wall stress. **A. Mkc1 phosphorylation during cell wall stress.** Cells of indicated genotypes were pre-grown in YPD medium with 5 ng/ml doxycycline for 3.5 h, and inoculated into YPD medium with 5 ng/ml doxycycline and 10 ng/ml micafungin (Mica). Protein extract was probed with antibody to phosphorylated Mkc1 (P-Mkc1) and the PSTAIRE antigen of Cdc28 as loading control. Dens: signal intensity ratio of P-Mkc1 to Cdc28. Representative of 2 biological replicates. Strains *TOR1/TOR1*, JKC1713; *tor1/TOR1*, JKC1347; *tetO-TOR1-Del381*, JKC1441; *tetO-TOR1,* JKC1549. **B. Growth during heat stress.** Dilutions of cells of indicated genotypes were spotted on YPD medium without or with 300 ng/ml doxycycline (300 Doxy) and 1M Sorbitol (Sor) and plates were incubated at 30° or 43°. Strains *TOR1/TOR1*, JKC1361; *tor1/TOR1*, JKC1347, JKC1346; *tetO-TOR1-Del381*, JKC1441, JKC1445; *tetO-TOR1*: *tor1/tetO-TOR1-FL,* JKC1549, JKC 1546. **C. Rps6 phosphorylation during heat stress.** Cells of indicated genotypes were pre-grown in YPD medium with 5 ng/ml doxycycline for 3.5 h, inoculated into YPD medium with 5 ng/ml doxycycline and incubated at 41° for indicated time periods. Total protein extract was probed with antibody to phosphorylated S6 (P-S6), and tubulin (Tub) as loading control. Dens: signal intensity ratio of P-S6 to Tub. Representative of 2 biological replicates. Strains *TOR1/TOR1*, JKC1713; *tor1/TOR1*, JKC1347; *tetO-TOR1-Del381*, JKC1441; *tetO-TOR1,* JKC1549.

**Fig. S5.**
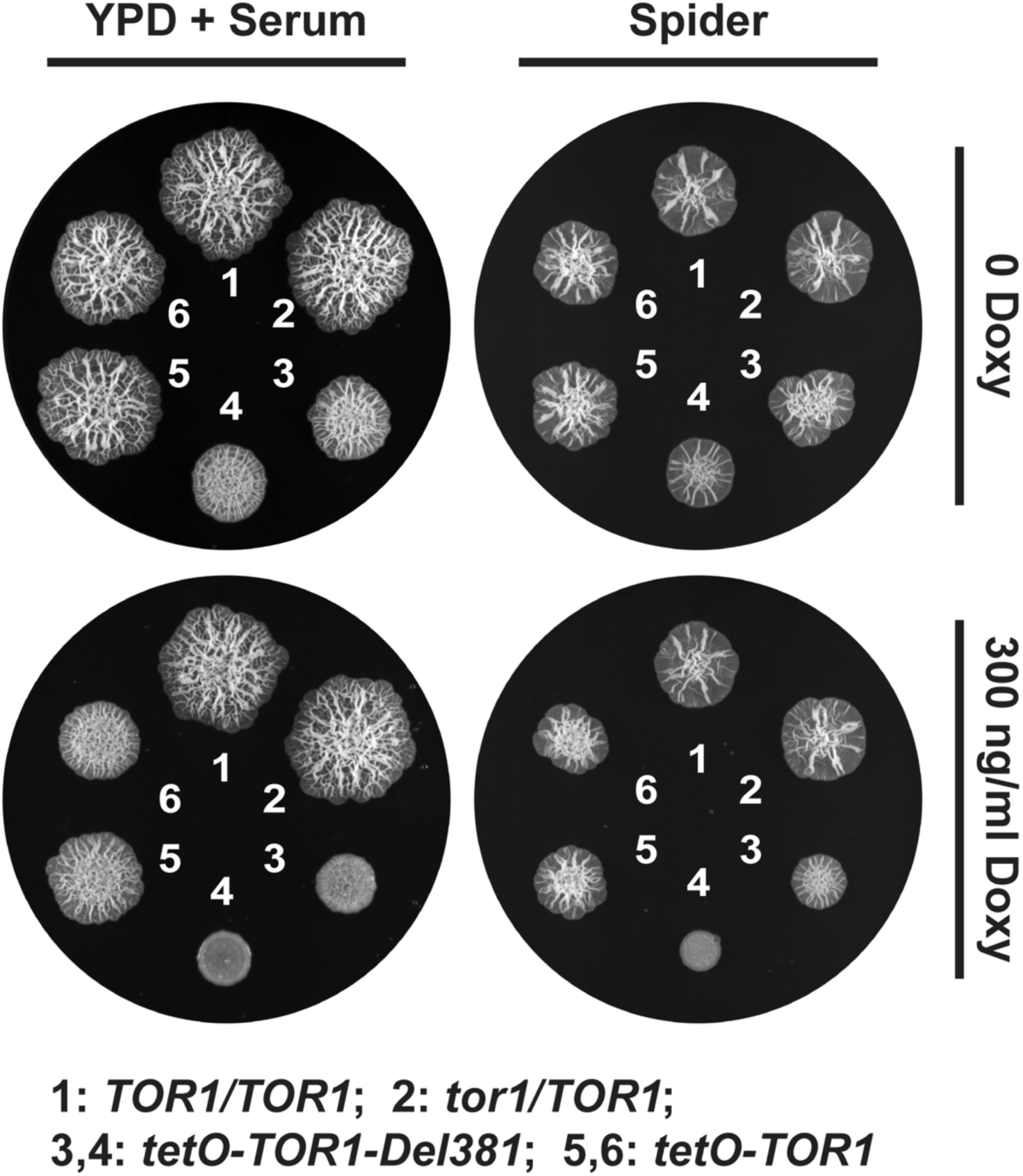
Spots of cells with distinct *TOR1* alleles showed specific surface wrinkling phenotypes. Cells of indicated genotypes were spotted at equidistant points around agar media, YPD+10% Serum or Spider medium (supplemented with 0.3 mM Histidine, and incubated at 37° for 3 days, without and with 300 ng/ml Doxycycline. Whole plates were imaged. Strains*TOR1/TOR1*, JKC1713; *tor1/TOR1*, JKC1347; *tetO-TOR1-Del381*, JKC1441, JKC1445; *tetO-TOR1,* JKC1549, JKC 1546.

**Table S1.** C. albicans strains used in this study.

| *C. albicans* strain name | Parent | Genotype | Strain background /construction | Reference |
| --- | --- | --- | --- | --- |
| SC5314 |  | Wild type |  | [1] |
| SN95 |  | *arg4Δ/arg4Δ his1Δ/his1Δ IRO1/iro1Δ::λimm<sup>434</sup> URA3/ura3Δ::λimm<sup>434</sup>* |  | [2] |
| JKC917 | SN95 | *hisΔ1/his1Δ::tetR-FRT arg4/arg4 IRO1/iro1Δ::λimm<sup>434</sup> URA3/ura3Δ::λimm<sup>434</sup>* |  | [3] |
| JKC1361 | JKC917 | *hisΔ1/his1Δ::tetR-FRT arg4/ARG4 IRO1/iro1Δ::λimm<sup>434</sup> URA3/ura3Δ::λimm<sup>434</sup>* | JKC917 transformed with PCR product of *fjk1184* & *rjk1186* using Candida cDNA library as template | [3] |
| JKC1713 | JKC917 | *hisΔ1/his1Δ::tetR-FRT arg4/ARG4 IRO1/iro1Δ::λimm<sup>434</sup> URA3/ura3Δ::λimm<sup>434</sup>* | JKC917 transformed with PCR product of *fjk1184* & *rjk1186* using Candida cDNA library as template | This work |
| JKC1347 | JKC917 | *tor1::ARG4/TOR1 hisΔ1/his1Δ::tetR-FRT arg4/arg4 IRO1/iro1Δ::λimm<sup>434</sup> URA3/ura3Δ::λimm<sup>434</sup>* | JKC917 transformed with PCR product of *fjk1142* & *rjk1143* using pFA-ARG4 as template | [3] |
| JKC1345 | JKC917 | *tor1::ARG4/TOR1 hisΔ1/his1Δ::tetR-FRT arg4/arg4 IRO1/iro1Δ::λimm<sup>434</sup> URA3/ura3Δ::λimm<sup>434</sup>* | *tor1/TOR1* heterozygote constructed like JKC1347, independent isolate. | This work |
| JKC1346 | JKC917 | *tor1::ARG4/TOR1 hisΔ1/his1Δ::tetR-FRT arg4/arg4 IRO1/iro1Δ::λimm<sup>434</sup> URA3/ura3Δ::λimm<sup>434</sup>* | *tor1/TOR1* heterozygote constructed like JKC1347, independent isolate. | This work |
| JKC1441 | JKC1347 | *tor1::ARG4/tor1::FRT-tetO-TOR1-Del381 hisΔ1/his1Δ::tetR-FRT arg4/arg4 IRO1/iro1Δ::λimm<sup>434</sup> URA3/ura3Δ::λimm<sup>434</sup>* | JKC1347 transformed with *StuI/NcoI* digested pJK1189 to have *TOR1-Del381* under *tetO (OFF)* promoter, after inducing FLP | [3] |
| JKC1442 | JKC1345 | *tor1::ARG4/tor1::FRT-tetO-TOR1-Del381 hisΔ1/his1Δ::tetR-FRT arg4/arg4 IRO1/iro1Δ::λimm<sup>434</sup> URA3/ura3Δ::λimm<sup>434</sup>* | *tor1/tetO-TOR1-Del381* constructed like JKC1441, distinct lineage from JKC1345. | This work |
| JKC1445 | JKC1346 | *tor1::ARG4/tor1::FRT-tetO-TOR1-Del381 hisΔ1/his1Δ::tetR-FRT arg4/arg4 IRO1/iro1Δ::λimm<sup>434</sup> URA3/ura3Δ::λimm<sup>434</sup>* | *tor1/tetO-TOR1-Del381* constructed like JKC1441, distinct lineage from JKC1346. | This work |
| JKC1549 | JKC1347 | *tor1::ARG4/tor1::FRT-tetO-TOR1-FL hisΔ1/his1Δ::tetR-FRT arg4/arg4 IRO1/iro1Δ::λimm<sup>434</sup> URA3/ura3Δ::λimm<sup>434</sup>* | JKC1347 transformed with *StuI/NcoI* digested pJK1236 to have *TOR1-FL* under *tetO (OFF)* promoter, after inducing FLP | [3] |
| JKC1543 | JKC1345 | *tor1::ARG4/tor1::FRT-tetO-TOR1-FL hisΔ1/his1Δ::tetR-FRT arg4/arg4 IRO1/iro1Δ::λimm<sup>434</sup> URA3/ura3Δ::λimm<sup>434</sup>* | *tor1/tetO-TOR1-FL* constructed like JKC1549, distinct lineage from JKC1345. | This work |
| JKC1546 | JKC1346 | *tor1::ARG4/tor1::FRT-tetO-TOR1-FL hisΔ1/his1Δ::tetR-FRT arg4/arg4 IRO1/iro1Δ::λimm<sup>434</sup> URA3/ura3Δ::λimm<sup>434</sup>* | *tor1/tetO-TOR1-FL* constructed like JKC1549, distinct lineage from JKC1346. | This work |
| TETG25B | CAI4 | *ADH1/adh1::tetO(ON)-GFP* | CAI4 transformed with pTET25 tetracycline inducible GFP cassette | [4] |
| JKC2616 | JKC1713 | *hisΔ1/his1Δ::tetR-FRT arg4/ARG4 MAL2/pMAL2-GFP-FRT-pMAL2-MAL2 IRO1/iro1Δ::λimm<sup>434</sup> URA3/ura3Δ::λimm<sup>434</sup>* | JKC1713 transformed with *BsrGI* digested pJK1489 to have GFP under *pMAL2* promoter, after inducing FLP | This work |
| JKC2620 | JKC1347 | *tor1::ARG4/tor1::FRT-tetO-TOR1-Del381 hisΔ1/his1Δ::tetR-FRT arg4/arg4 MAL2/pMAL2-GFP-FRT-pMAL2-MAL2 IRO1/iro1Δ::λimm<sup>434</sup> URA3/ura3Δ::λimm<sup>434</sup>* | JKC1347 transformed with *BsrGI* digested pJK1489 to have GFP under *pMAL2* promoter, after inducing FLP | This work |
| JKC2624 | JKC1441 | *tor1::ARG4/tor1::FRT-tetO-TOR1-Del381 hisΔ1/his1Δ::tetR-FRT arg4/arg4 MAL2/pMAL2-GFP-FRT-pMAL2-MAL2 IRO1/iro1Δ::λimm<sup>434</sup> URA3/ura3Δ::λimm<sup>434</sup>* | JKC1441 transformed with *BsrGI* digested pJK1489 to have GFP under *pMAL2* promoter, after inducing FLP | This work |
| JKC2628 | JKC1549 | *tor1::ARG4/tor1::FRT-tetO-TOR1-FL hisΔ1/his1Δ::tetR-FRT arg4/arg4 MAL2/pMAL2-GFP-FRT-pMAL2-MAL2 IRO1/iro1Δ::λimm<sup>434</sup> URA3/ura3Δ::λimm<sup>434</sup>* | JKC1549 transformed with *BsrGI* digested pJK1489 to have GFP under *pMAL2* promoter, after inducing FLP | This work |

**Table S2.**
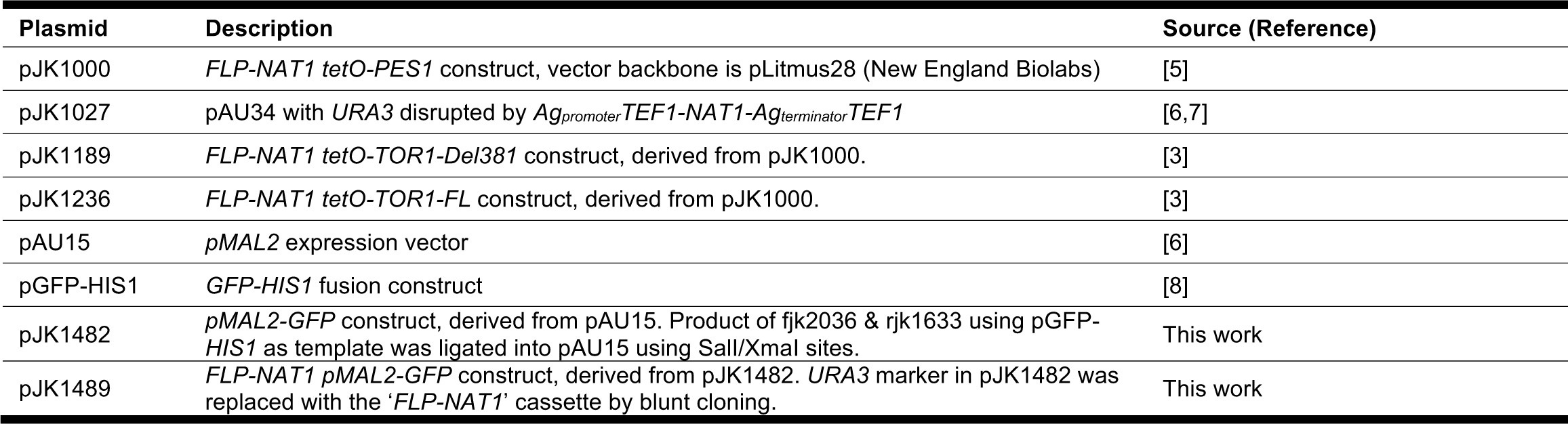
Plasmids used in this study.

**Table S3.**
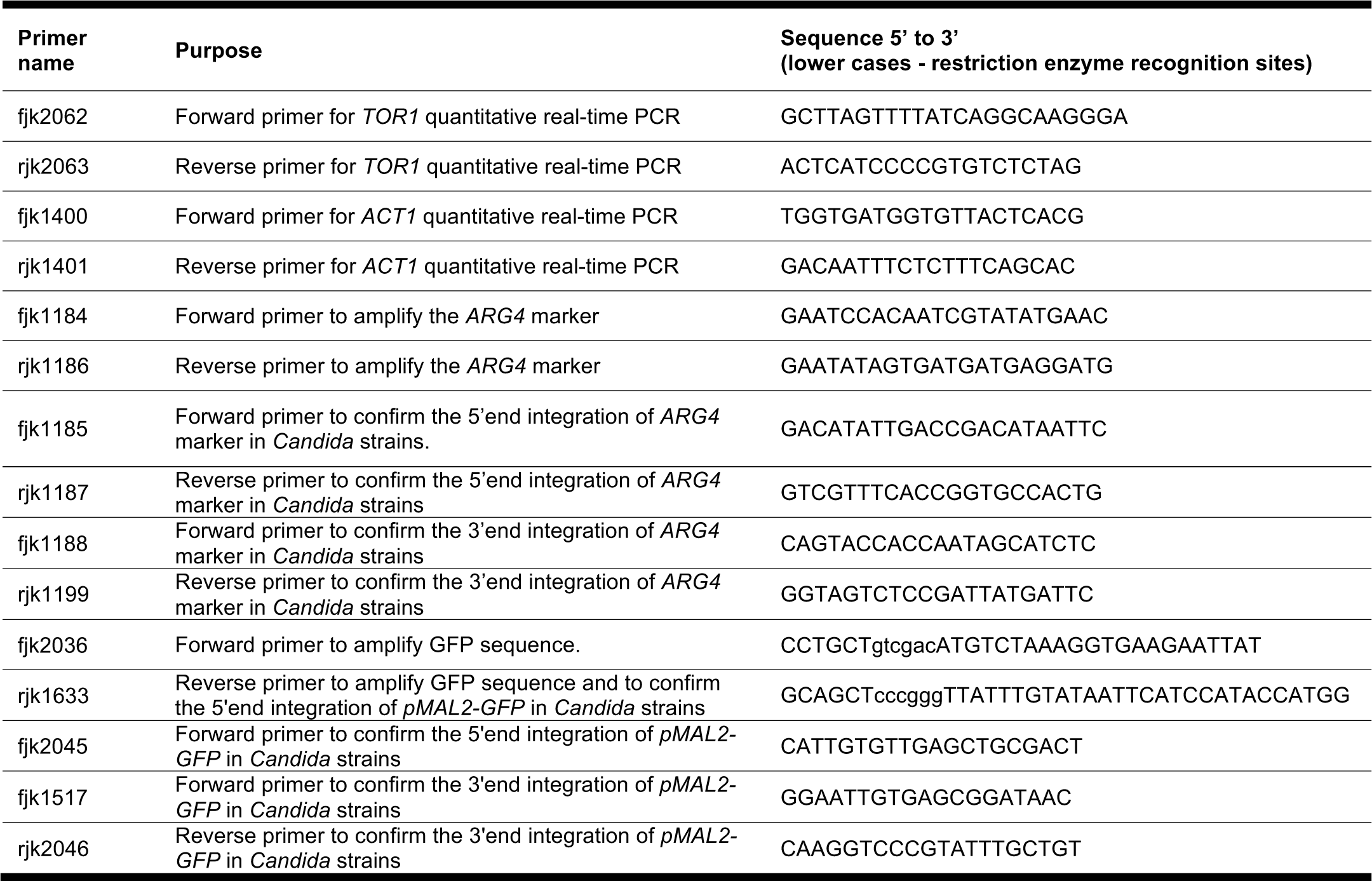
Oligonucleotides used in this study.

**Table S4.**
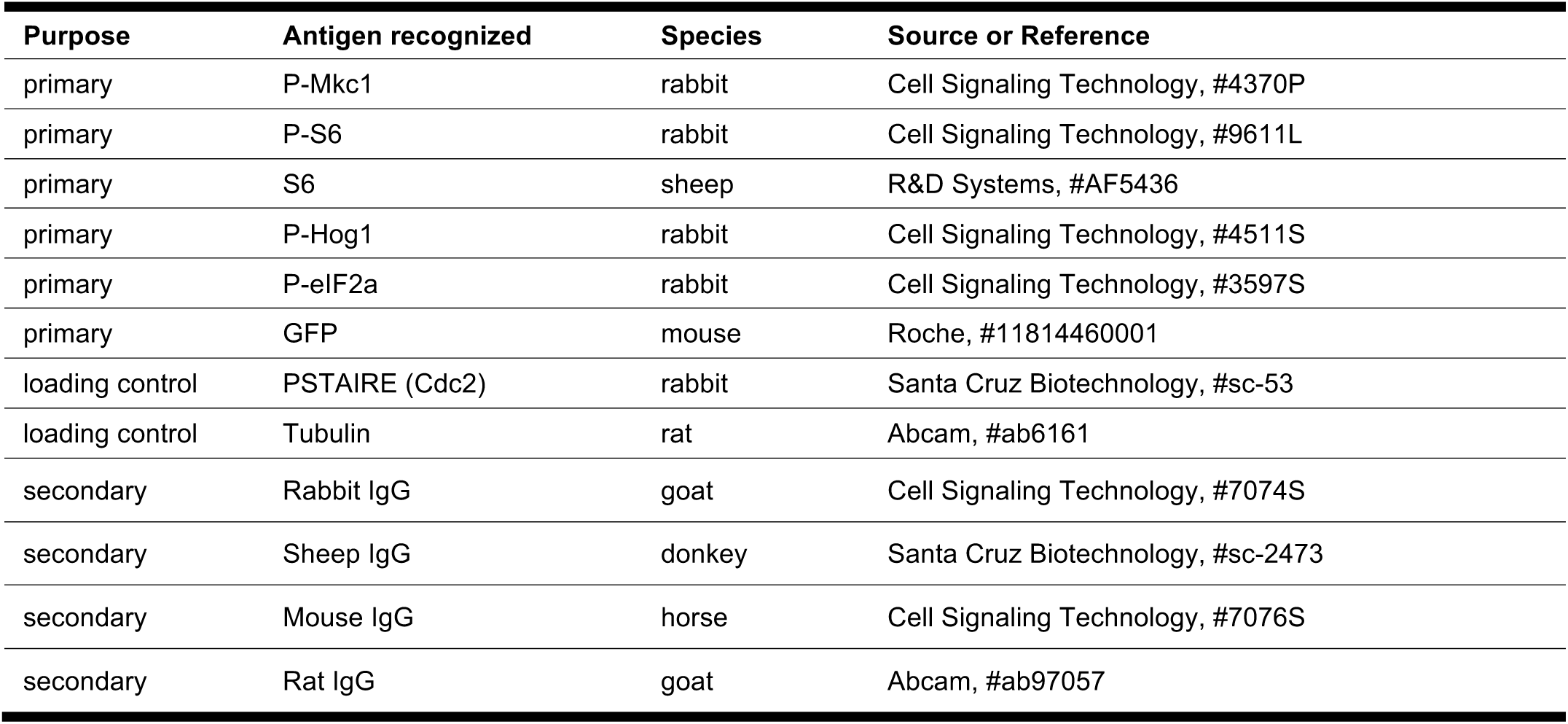
Antibodies used in this study.

